# Noradrenergic responsiveness supports selective attention across the adult lifespan

**DOI:** 10.1101/551879

**Authors:** Martin J. Dahl, Mara Mather, Myriam C. Sander, Markus Werkle-Bergner

## Abstract

Selectively attending to relevant information while blocking out distractors is crucial for goal-directed behavior, yet with advancing age, deficits emerge in attentional selectivity. Decrements in attention have been associated with altered noradrenergic activity in animals. However, research linking noradrenergic functioning to attention in aging humans is scarce, likely reflecting long-standing methodological challenges in non-invasive assessments. We studied whether age-related differences in the noradrenergic system predict differences in attention. We measured pupil dilation, a non-invasive marker of phasic norepinephrine (NE) release, while concurrently recording the electroencephalogram (EEG), of male younger (N=39; 25.2±3.2 years) and older adults (N=38; 70.6±2.7 years). Arousal was triggered on a trial-by-trial basis using fear-conditioned (CS+) stimuli. During conditioning, pupil and EEG markers related to heightened NE activity were identified. Afterwards, in a dichotic listening task, participants were cued to direct attention to either the left or right ear while highly similar syllable pairs were presented simultaneously to both ears. During the dichotic listening task, presentation of fear-conditioned stimuli reinstated the acquired fear response, as reflected in pupil and EEG alpha–beta-band responses. Critically, pupil dilation to CS+ was correlated with stronger EEG alpha–beta desynchronization, suggesting a common dependence on NE release. On a behavioral level, stronger arousal reactions were associated with better attention. In particular, structural equation modeling revealed that the responsiveness of the NE system is associated with attention on a latent construct level, measured by several indicator tasks. Overall, our results suggest that the responsiveness of the NE system supports attention across the lifespan.

**Significance statement:** In old age the ability to selectively process relevant aspects of the environment fades. Animal research suggests that the neuromodulator norepinephrine helps to maintain selective attention. We tested younger and older adults across a variety of attention tasks. In addition, we used arousing stimuli to experimentally activate participants’ noradrenergic system while recording pupillometry and electroencephalography (EEG) to infer its functional capacity. Older adults showed compromised attention and reduced noradrenergic responsiveness as indicated by interrelated pupil and EEG markers. Crucially, in both age groups a more responsive noradrenergic system was strongly associated to attention. Our findings link animal and human studies on the neural underpinning of attention in aging and underscore the importance of the noradrenergic system in late life cognition.

## Introduction

Daily situations confront us with a plethora of competing sensory inputs that far exceed neural processing capacities, thus prioritization and selection is essential for adaptive behavior (e.g., Desimone & Duncan, 1995). Impaired attentional selection in aging (for reviews see Kennedy & Mather, 2019; Plude, Enns, & Brodeur, 1994) has been linked to deficient neuromodulation (Li et al., 2001; Bäckman et al., 2006). The neuromodulator norepinephrine (NE) is strongly implicated in attentional processes that facilitate the processing of relevant information (Berridge & Waterhouse, 2003). First, increased NE release is associated with the transition to and the maintenance of an activated cortical and behavioral state – as evident in a desynchronized (high frequency, low amplitude) electroencephalogram (EEG) and alert waking (sometimes termed arousal; Carter et al., 2010; McGinley et al., 2015; Neves et al., 2018). In the waking state, fast, burst-like (phasic) and slow, rhythmic (tonic) firing patterns of the locus coeruleus (LC), the primary cortical NE source, have been tied to focused attention and distractibility, respectively (Aston-Jones and Cohen, 2005). Further, a series of pharmacological and lesion studies demonstrated that, via actions at α_2A_-adrenoceptors in the prefrontal cortex, NE facilitates top-down selective attention (Arnsten & Li, 2005). In the sensory cortices, phasic NE release interacts with local glutamate levels to allow the selective processing of currently relevant representations, mediated via α_2A_- and ß-adrenoceptors (Mather et al., 2016; Gelbard-Sagiv et al., 2018). Finally, NE has been linked to the reorienting and switching of attention via disruption of the dorsal- and activation of the ventral attention network (Bouret & Sara, 2005; Corbetta, Patel, & Shulman, 2008). In line with these links between NE and attention, recent theories of both healthy (Mather and Harley, 2016) and pathological (Weinshenker, 2018; Satoh and Iijima, 2019) cognitive aging have proposed a prominent role of the LC-NE system in late life cognition. However, LC’s anatomical location in the brainstem, adjacent to the ventricular system and its widespread, unmyelinated axons expose it to blood- and cerebrospinal-fluid bound toxins, making it vulnerable to neurodegeneration (Mather and Harley, 2016; Liu et al., 2019) with potentially wide-ranging consequences. For instance, Wilson and colleagues (2013) reported an association between LC’s structural integrity, as assessed post-mortem via autopsy, and longitudinal cognitive decline in aging (cf. Hämmerer et al., 2018; Betts et al., 2019; Dahl et al., 2019b). However, the question of how LC’s functional characteristics, i.e., its capacity to respond to behaviorally relevant information, are linked to attention in aging humans is still unresolved. Long-standing technical challenges in the non-invasive assessment of LC-NE activity in vivo (e.g., Astafiev, Snyder, Shulman, & Corbetta, 2010) have contributed to this lack of information. However, recently, multiple independent studies (Joshi et al., 2016; Reimer et al., 2016; Breton-Provencher and Sur, 2019; Zerbi et al., 2019) demonstrated that pupil dilation in the absence of interfering visual input serves as valid, non-invasive proxy for LC activity. In addition, use of optogenetics established a causal link between phasic LC activity and event-related EEG responses (i.e., P300 ERP; Vazey, Moorman, & Aston-Jones, 2018). Moreover, EEG reveals fluctuations in cortical states (i.e., global EEG de/activation as reflected in a de/synchronized EEG) that have been associated with LC activity (McCormick et al., 1991).

In this study we thus used a multimodal assessment to evaluate individual differences in selective attention among younger and older adults and their dependence on functional characteristics of the LC-NE system. In order to experimentally induce LC activity, we made use of LC-NE’s well-established role in fear processing (cf. Lee et al., 2018; Szabadi, 2012; Uematsu et al., 2017). We hypothesized that the functional capacity of the LC-NE system as assessed by simultaneous, interrelated pupil and EEG responses would be closely associated with individual differences in selective attention. In sum, the overall goal of this study was to extend our knowledge about the role of the LC-NE system in human cognitive aging by generating a multimodal, non-invasive index of LC functioning and linking it to attention abilities in younger and older adults.

## Methods

### Study design

Data was collected within the framework of a larger study investigating the interplay of neurophysiological indices of LC-NE activity and their association to selective attention in younger and older adults (YA; OA, respectively). Only aspects of the study that are relevant to the current analyses are introduced in detail below.

Participants were invited on three successive days (Day 1–Day 3) for individual assessments that spanned approximately 4 hours on Day 1 and Day 2 and 2 hours on Day 3. Time of assessment (morning, afternoon, evening) was kept constant across sessions within participants.

In short, on the first day, participants completed a neuropsychological selective attention battery as well as an assessment of fear conditionability (see Figure 1 a and c) while pupil dilation was recorded. To adapt auditory stimuli during later attention assessments for hearing thresholds, we assessed hearing acuity (on Day 1 for younger adults and on a separate occasion preceding Day 1 for older adults). On the second day, we concurrently recorded pupil dilation and EEG while participants underwent another fear conditioning session and an in-depth evaluation of their auditory selective attention performance (see Figure 1b). The last day comprised a final fear conditioning session while recording pupil dilation (see Figure 1c) and an MRI assessment which is reported in more detail here: https://doi.org/10.17605/OSF.IO/G9FQJ. The study was approved by the ethics committee of the German Psychological Association (DGPs) and was conducted in accordance with relevant guidelines and regulations.

**Figure 1.**
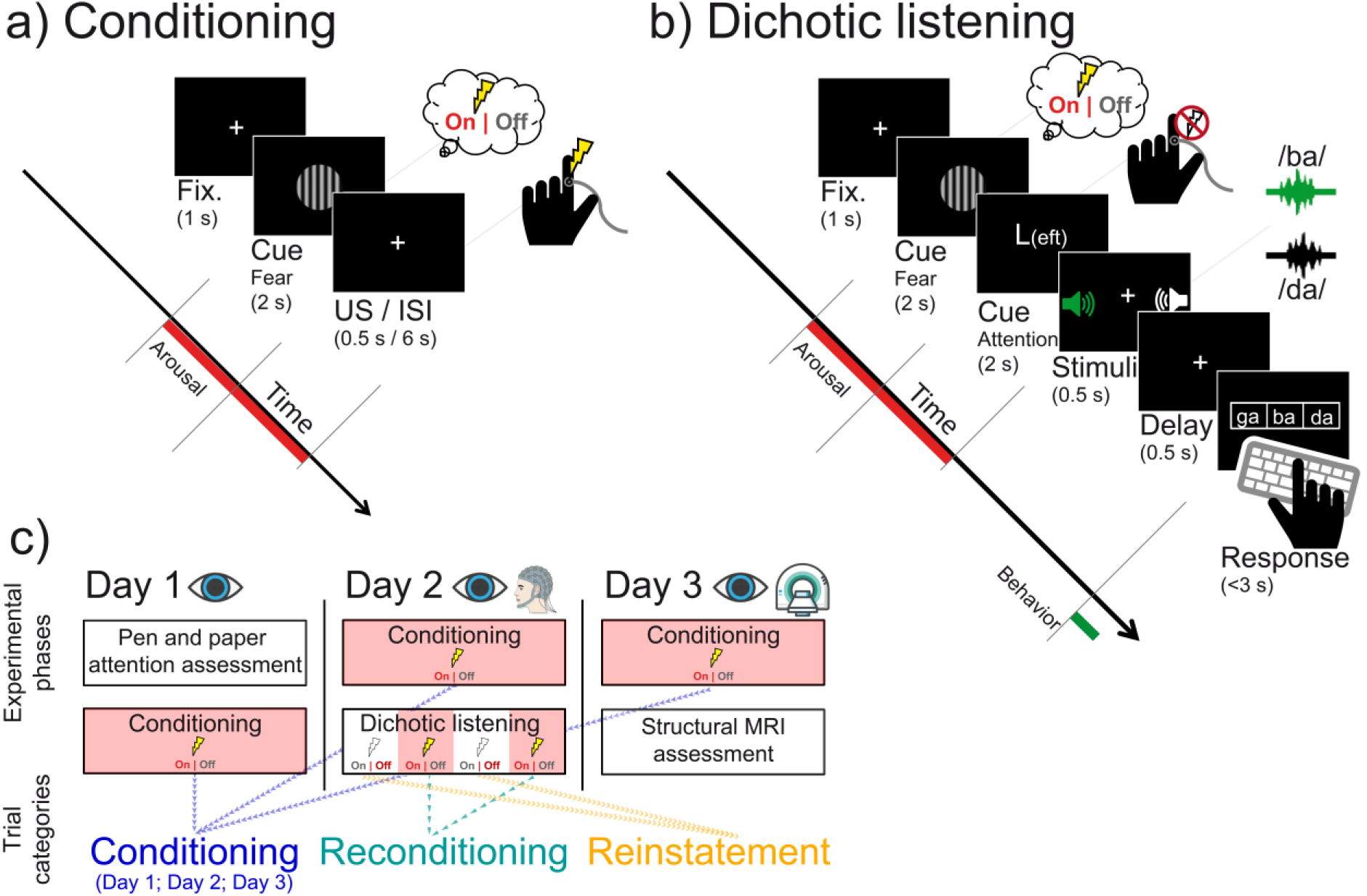
Experimental procedure of the (a) fear conditioning session and (b) dichotic listening task as well as (c) resulting trial categories. (a) Each conditioning trial began with a 1 s baseline interval during which a white fixation cross was presented. Then, either the conditioned (CS+) or perceptually matched control stimulus (CS–; horizontal or vertical Gabor patch) was displayed for 2 s. In CS+ trials, upon offset of the Gabor patch, a mild electric shock was applied to participant’s finger. After 6 s of fix cross presentation (inter-trial-interval), the next trial started. (b) Importantly, the beginning of trials in the dichotic listening task was identical to the fear conditioning session. Upon offset of the CS, an attention cue (2 s) indicated which ear participants should focus on. Tightly synchronized syllable-pairs were then presented simultaneously to the left and right ear (0.5 s) and participants indicated by button press which syllable they heard on the cued ear (up to 3 s). (c) Participants underwent fear conditioning sessions (cf. a) on each day of the experiment while pupil dilation was recorded (Conditioning; blue). The arousal manipulated dichotic listening task (cf. b) was completed once on the second day while both pupil dilation and EEG was recorded. Interleaved after each break of the dichotic listening task participants were reconditioned (cf. a) to prevent extinction of the fear response (Reconditioning; teal). During re/conditioning participants received electrical stimulation (US) and thus the observed responses may represent a mixture of fear and somatosensory responses. In contrast, during the dichotic listening task no shocks were applied and thus observed responses to the arousal manipulation (CS+ vs CS–) indicate the reinstatement of the fear response (Reinstatement; orange). On the first day of the experiment, participants additionally completed a pen and paper attention assessment while on the last day structural magnetic resonance imaging (MRI) data was recorded. Fix.: Fixation; US: Unconditioned stimulus; ISI: Inter stimulus interval. Numbers in brackets indicate presentation times in seconds (s).

### Participants

Forty-one younger adults participated in the study. Two of these (4.88%; aged: 24.59 and 29.02 years) were not re-invited after Day 1 due to low quality eye tracking data, reducing the final sample to 39 younger adults (mean (*M*) age: 25.23 ± 3.23 years (standard deviation; *SD*); range: 20.17–31 years; 100% male). In addition, thirty-eight older adults took part in the experiments (*M* age: 70.61 ± 2.71 years (*SD*); range: 65.50–75.92 years; 100 % male). All participants were healthy, MRI-compatible, right-handed, fluent German speakers with normal or corrected-to-normal vision who provided written informed consent and were reimbursed for participation. Intake of centrally active drugs and in particular medication directly influencing the LC-NE system (e.g., beta blockers) precluded participation. Please note that the current study only tested male subjects due to our pilot data demonstrating reliable sex differences in the stimulation intensity participants selected for the fear conditioning sessions. Previous research indicates sex differences in the capability to learn and maintain fear responses during conditioning (Merz et al., 2018; but also see Gruene et al., 2015 and Voulo and Parsons, 2017, for evidence for sex-specific expressions of fear learning). Some of these differences in fear learning may be associated with sex differences in the LC-NE system (Bangasser et al., 2016; Mulvey et al., 2018). To limit the number of control variables, we decided to test male participants; however, we would like to underscore the need to follow up on the reported findings including both sexes. Descriptive characteristics of the two age groups are provided in Table 1. Both groups showed comparable educational levels and did not differ reliably on a brief dementia screening (Mini Mental State Examination (MMSE); Folstein, Folstein, & McHugh, 1975). All participants scored above the commonly used MMSE cut-off of 26 points. In line with previous reports (e.g., Passow et al., 2012, 2014), older adults demonstrated higher scores on a test of verbal knowledge (Spot a Word; Lehrl, 1977) and increased age-related hearing loss. On average, older adults selected higher intensities as unpleasant unconditioned stimulus for the fear conditioning procedure, presumably reflecting age related differences in skin conductivity (Chamberlin et al., 2011).

**Table 1.** Descriptive statistics for younger and older adults

| Measure | Younger adults<br>(n = 39) | Older adults<br>(n = 38) | *U* | *Z* | *p* |
| --- | --- | --- | --- | --- | --- |
| Age (years) | 25.231 ± 0.516 | 70.614 ± 0.440 | 780 | -7.546 | <0.001 |
| Education (years) | 16.171 ± 0.406 | 17.263 ± 0.560 | 1320 | -1.487 | 0.137 |
| Hearing level (dB) | 8.027 ± 0.758 | 19.626 ± 1.075 | 907 | -6.251 | <0.001 |
| Shock intensity (mA) | 0.3 ± 0.026 | 0.4 ± 0.024 | 1211 | -3.227 | 0.001 |
| Spot-A-Word (correct rows) | 21.487 ± 0.551 | 29.158 ± 0.382 | 834 | -7.021 | <0.001 |
| Mini Mental State (points) | 29.410 ± 0.120 | 29.053 ± 0.181 | 1631 | 1.216 | 0.224 |

Note: For younger and older adults, group means ± 1 standard error of the mean are reported. Age group comparisons are based on non-parametric Mann-Whitney U-tests. Hearing level is averaged across frequencies (250–3000 Hz) and ears. Shock intensity denotes the intensity participants individually defined as uncomfortable yet not painful on Day 2.

### Experimental procedures and stimuli

#### Neuropsychological attention assessment

On the first day of the experiment, participants completed a multimodal, standardized neuropsychological attention assessment comprising the D2 test of attention (Brickenkamp and Zillmer, 1998), Digit-Symbol-Substitution Test (Wechsler, 1981) and auditory digit sorting task (cf. Kray & Lindenberger, 2000).

The D2 test is a paper and pencil cancellation task asking participants to cross out any letter *d* with two marks (‘’) around, above or below it from a stream of highly similar surrounding distractors (e.g., *p* with two marks or *d* with only one mark). Participants were granted 20 s to complete each of a total of 12 lists of items. The difference between processed items and committed errors (errors of omission and errors of commission) across lists was taken as a measure of attention.

During the Digit-Symbol-Substitution Test participants were presented with a list of digit-symbol pairs (e.g., 1: –; 2: ⏊; […] 9: =) along with a list of digits. Participants were asked to write down the corresponding symbol under each digit as quickly and accurately as possible. The number of correctly copied symbols within 90 s was taken as an index of attention. In the auditory digit sorting task we auditorily presented participants with a stream of numbers (e.g., 2–7–5) ranging from three to eight digits. Participants then had to write down the numbers sorted according to numerosity (e.g., 2–5–7). The sum of correctly reported answers across all trials was taken as a measure of attention.

#### Fear conditioning

To experimentally activate the LC-NE system (cf. Rasmussen & Jacobs, 1986; Szabadi, 2012; Uematsu et al., 2017) on each assessment day (Day 1–3), participants completed a brief fear conditioning session in which they learned the association between a visual stimulus and an aversive electrical shock (cf. Lee et al., 2018; see Figure 1a). During this phase, either a horizontal or vertical sinusoidal luminance pattern (i.e., Gabor patch; CS+) was paired with an unconditioned stimulus (US; shock). The other pattern was never paired with the US and served as perceptually matched control stimulus (CS–); the association between pattern orientation (horizontal/vertical) and shock was kept constant within subject across days and was counterbalanced within age groups (YA: 21:18; OA: 20:18). Note that this design guaranteed a matched luminance of CS+ and CS– while the former acquired the behavioral relevance to stimulate LC-NE activity (Szabadi, 2012). Each conditioning session comprised 40 trials which started with a central white fixation cross on a black background (baseline; 1 s), followed by the visual stimulus (2 s; CS+ or CS–; see Figure 1a). After offset of the visual stimulus the shock was applied in CS+ trials for 0.5 s with a 80% reinforcement schedule (i.e., 0 s trace conditioning) using a ring electrode connected to a bipolar current stimulator (DS5; Digimeter; Welwyn Garden City, United Kingdom) that was taped either to participant’s left or right index finger (hand assignment was constant within subject across days and counterbalanced within age groups: YA: 19:20; OA: 19:19). The inter-trial-interval (ITI; white fixation cross) was set to 6 s to allow sufficient time for the pupil to return to baseline in CS+ trials (cf. Lee et al., 2018). The conditioning phase consisted of 20 CS+ and 20 CS– trials that were presented in pseudorandomized order. After half of the trials, participants had a self-paced break during which they were asked to indicate which of the two visual stimuli (horizontal; vertical) was paired with the shock and how many shocks were delivered (i.e., manipulation check). On each day, prior to the experiment participants individually selected an intensity for the US which they perceived as unpleasant but not painful (cf. Lee et al., 2018; see Table 1). During all conditioning sessions, gaze position and pupil dilation was recorded, interfering sensory input was minimized and central fixation was enforced (see below).

During data collection, for one older adult, assignment of Gabor patch orientation (horizontal/vertical) to CS condition (CS–/CS+) was switched between the first and second day of assessment. Since this error worked against finding a reliable difference in responses to CS– and CS+, we decided not to exclude this subject.

#### Dichotic listening task

On the second assessment day, we probed selective auditory attention by cueing participants to focus attention to either the left (focus left condition; FL) or right (focus right condition, FR) ear while highly similar consonant-vowel (CV) syllable pairs were presented dichotically (i.e., simultaneously one stimulus to the left and one to the right ear). Only syllables played to the cued ear should be reported while distractor stimuli should be ignored. To indicate their response, after a brief delay participants selected the target syllable from three visually displayed response options (including the target, distractor and one highly similar novel, i.e., not presented, syllable; see Figure 1b).

Within attentional conditions (FL and FR) we manipulated participant’s arousal level on a trial-by-trial basis. In particular, each trial started with a central white fixation cross on a black background (baseline; 1 s), followed by one of the conditioned stimuli (2 s; CS+ or CS–). After offset of the CS, a letter was centrally displayed cuing participants to adapt their attentional focus (2 s; *L* for FL; *R* for FR). Next, a syllable pair was presented dichotically for 0.5 s. After a delay of 0.5 s a recognition matrix (containing the target, distractor and one novel syllable) was visually displayed for up to 3 s and participants indicated by button press which syllable they heard in the cued ear. Response hand assignment was counterbalanced within age groups (YA: 20:19; OA: 19:19; shocks were never applied to the response hand). The ITI was set to 0.5 s and consisted of a white fixation cross. Matched trial timing between fear conditioning and dichotic listening tasks (1 s baseline; 2 s CS; ∼6 s until next trial) allows a comparison of arousal responses across both tasks (see Figure 1).

Twelve consonant-vowel syllable pairs consisting of syllables of voiced (/b/, /d/, /g/) or unvoiced (/p/, /t/, /k/) consonants combined with the vowel /a/ served as auditory stimuli in the dichotic listening task. Each pair contained two syllables with the same voicing that were matched for onset times (cf. Hugdahl et al., 2009; Westerhausen et al., 2009). To account for age-related hearing loss, syllable pairs were presented at an individually adjusted volume (i.e., 65 dB above participant’s average hearing threshold between 250 and 3000 Hz as assessed by means of pure-tone audiometry; cf. Passow et al., 2014). All twelve dichotic syllable pairs were presented six times in each of the attention and arousal conditions, summing to 288 trials in total (i.e., 12 syllable pairs × 6 presentations × 2 attentional focus × 2 arousal conditions) that were split in blocks of 48 trials. In 8.33% of the trials, no conditioned stimulus (CS+; CS–) was displayed (no-CS trials; n = 24), with a fixation cross instead occurring at that time point to obtain an index of auditory attention irrespective of CS presentation. In another 8.33% of the trials, the CS+ was followed by an electrical shock (booster trials; cf. fear conditioning phase; n = 24; cf. Lee et al., 2018) to prevent extinction of the conditioned response. No-CS and booster trials were excluded from analyses. After each block, participants had a self-paced break during which their average accuracy was displayed graphically. Breaks were followed by a brief reconditioning period of 20 trials that resembled one half of the fear conditioning phase (presented in pseudorandomized order) to maintain the fear response throughout the experiment (see Figure 1c).

To thoroughly familiarize participants with both the auditory material and the instructions prior to testing, on the first assessment day we presented the six syllables first diotically (i.e., the same syllable at the same time to the left and right ear; 24 trials) followed by a presentation to only one ear (6 left and 6 right ear trials). Participants indicated by button press which syllable they heard / on which ear (mean accuracy: 94.801 ± 0.983 (SEM) %). In addition, participants completed a dichotic listening task without arousal manipulation (96 trials; mean accuracy: 46.749 ± 0.010 (SEM)%; please note that chance performance in this task is 33.3%, i.e., one of three possible choices).

All stimuli were presented using Psychtoolbox (<u>Psychophysics Toolbox</u>, RRID:SCR_002881) for Matlab (<u>MATLAB</u>, RRID:SCR_001622; The MathWorks Inc., Natick, MA, USA) and insert earphones (ER 3A; Etymotic Research, Inc. Elk Grove Village, IL, USA). During the fear conditioning and dichotic listening task, gaze position, pupil dilation and the EEG (Day 2 only) were continuously recorded (see below). To minimize the influence of sensory input on pupil dilation, testing took place in a dark, sound-attenuated and electro-magnetically shielded room (cf. Hong, Walz, & Sajda, 2014). Further, to minimize the influence of eye movements on pupil dilation (Gagl et al., 2011) at the beginning of each trial (baseline period) participant’s gaze position was sampled online and the trial only started if central fixation was either maintained (> 75% of the time) or restored upon presentation of a re-fixation target.

### Behavioral analyses

#### Dichotic listening task

To evaluate age differences in selective attention, we calculated the auditory laterality index (LI; Marshall, Caplan, & Holmes, 1975), for each Attentional focus condition (FL, FR), collapsing across arousal conditions. This index expresses the amount of right relative to left ear responses (i.e., LI = (Right – Left) / (Right + Left)). The LI ranges from –1 to 1 whereby negative values indicate more left ear responses and positive values index a tendency towards selecting the right ear syllable. Younger and older adults’ laterality indexes were analyzed in a two-factorial (Age group × Attentional focus (FL, FR)) mixed measures analysis of variance (ANOVA) that was followed-up by post-hoc tests within age groups. To judge the influence of age-related hearing loss on auditory selective attention, in a second ANOVA (Age group × Attentional focus) we included participants’ average hearing loss as a covariate. For further analyses, the difference between laterality indexes for the FL and FR condition was calculated to provide an overall measure of participant’s auditory selective attention ability.

#### General attention

To obtain a single measure for general attention performance, i.e., independent of the specific task used, we made use of the comprehensive cognitive battery available for this data set. In particular, we integrated performance across multiple visual and auditory attention tasks (i.e., dichotic listening task; D2 test of attention; digit-symbol-substitution task; auditory digit sorting task) by means of a structural equation modeling approach (SEM; see Figure 2, lower part) using the Ωnyx 1.0-1013 software package (von Oertzen et al., 2015) with full information maximum likelihood estimation (FIML). SEM offers a multivariate approach in which observed (manifest) variables can be used to examine hypotheses about unobserved (latent) variables. Latent variables have the benefit of accounting for measurement error in observed scores and thereby increasing statistical power (Curran et al., 2010; Kievit et al., 2018).

**Figure 2.**
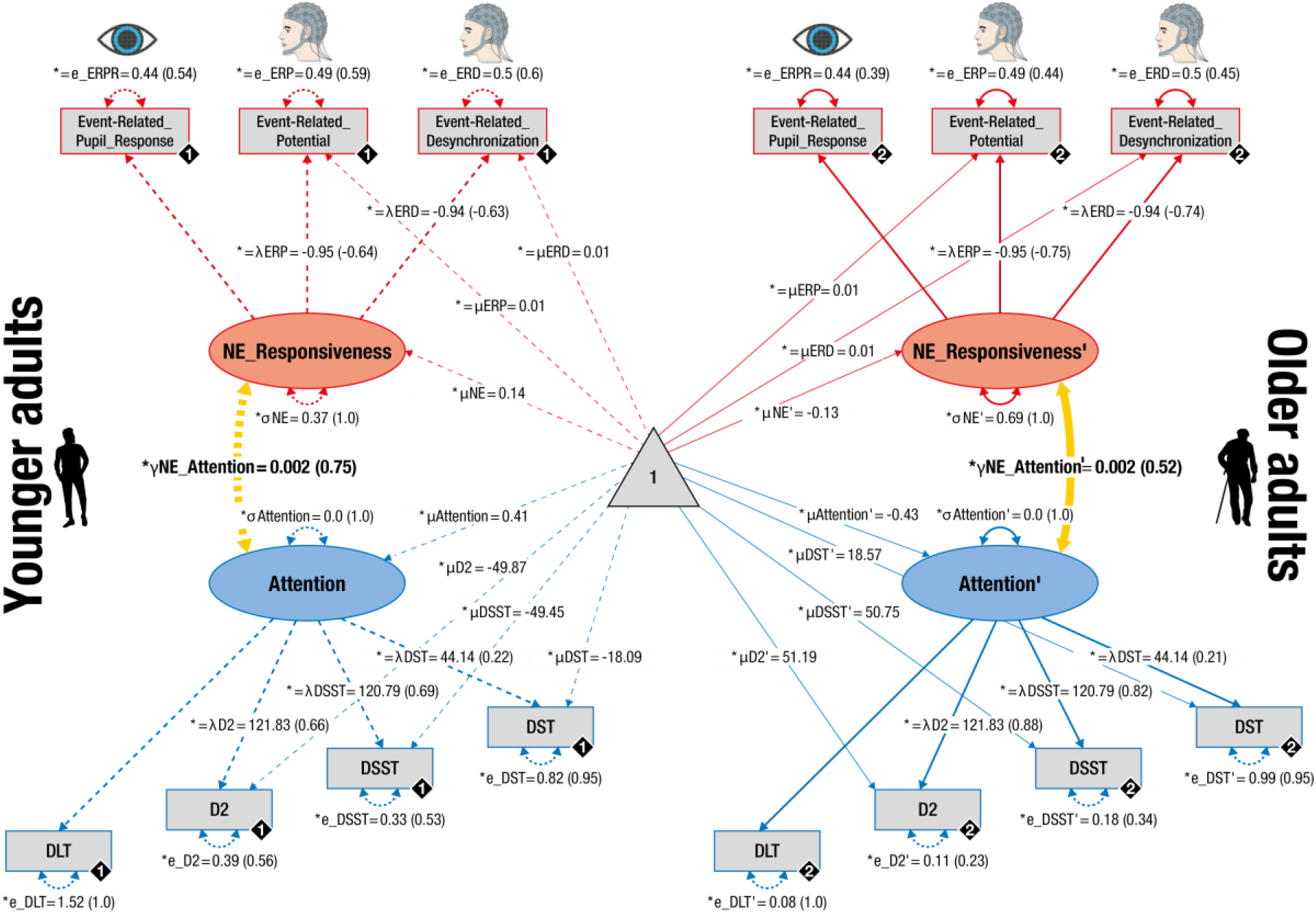
Pictorial rendition of the structural equation model that probes associations (yellow lines) between noradrenergic responsiveness (NE_Responsiveness; red) and attention performance (blue) in younger and older adults on a latent level. Rectangles and ellipses indicate manifest (observed) and latent variables, respectively. The constant is depicted by a triangle. Cognitive manifest variables represent attention performance assessed in a dichotic listening task (DLT; cf. Figure 1 and 3), D2 task of attention (D2; cf. Table 2), digit-symbol-substitution task (DSST) and digit sorting task (DST). Physiological manifest variables represent the reinstatement of fear conditioned Event-related Pupil Responses (ERPR; cf. Figure 4a, 5a), Event-related Potentials (ERP; cf. Figure 4a) and Event-related Desynchronization (ERD; cf. Figure 5a). Black diamonds on manifest variables indicate the age group (younger adults = 1, broken lines; older adults = 2, solid lines). (Co)Variances (γ, σ) and loadings (λ) in brackets indicate standardized estimates. Loadings that are freely estimated (*) but constrained to be equal across age groups (=) are indicated by both asterisk and equal signs (*=). Note that the cognitive sub-model demonstrated metric factorial invariance (invariant manifest means and errors across age groups) whereas the physiological sub-model showed strict factorial invariance (manifest means and errors are constrained across groups).

**Table 2.** Overview of age differences in attention

Overview of age differences in attention
| Task | N (YA/OA) | PerformanceYA<br>(z±SEM) | Performance OA<br>(z±SEM) | Z | p |
| --- | --- | --- | --- | --- | --- |
| DLT | 77 (39/38) | 0.415 (0.200) | -0.425 (0.048) | 3.790 | <0.001 |
| D2 | 77 (39/38) | 0.634 (0.133) | -0.651 (0.113) | 6.109 | <0.001 |
| DSST | 77 (39/38) | 0.629 (0.129) | -0.645 (0.120) | 5.689 | <0.001 |
| DST | 77 (39/38) | 0.206 (0.153) | -0.212 (0.164) | 2.066 | 0.039 |

Note: For each age group for all cognitive tasks mean performance is provided as z-value ± the standard error of the mean. DLT: Dichotic listening task; D2: D2 test of attention; DSST: Digit-Symbol-Substitution Task; DST: Digit-Sorting Task; YA: Younger adults; OA: Older adults; All comparisons are evaluated using non-parametric Mann-Whitney U-tests (YA vs. OA).

In particular, in each age group standardized performance in the dichotic listening task, D2 test of attention, digit-symbol-substitution and digit sorting tasks served as manifest variables and loaded on a single latent selective attention factor (i.e., a multiple-group model). Factor loadings (other than the first, which was fixed to one) were estimated freely but were constrained to be equal across groups. The model demonstrated metric factorial invariance (i.e., it required variant manifest intercepts and variances across groups) and thus precluded an interpretation of age group differences in the means of the latent factor (Meredith and Teresi, 2006; Schwab and Helm, 2015). We assessed the adequacy of the proposed selective attention model by testing for differences between the model-implied and empirically observed covariance matrices (Eid et al., 2015). The *χ*^2^-test formally tests for equity of the covariance matrices. However, since it is particularly sensitive to sample size it should be interpreted with caution in large samples (Brown, 2006; Eid et al., 2015). We thus additionally examined two frequently reported fit indices: First, the root mean square error of approximation (RMSEA) that is a closeness of fit coefficient expressing how much the postulated model approaches the true model. Second, the comparative fit index (CFI), an incremental fit index which compares the goodness of fit of the proposed model with a more restrictive nested baseline model (Brown, 2006; Curran et al., 2010; Eid et al., 2015). In contrast to the *χ*^2^-test, the RMSEA and CFI are not influenced by sample size. RMSEA values close to or below 0.06 and CFI values of close to 0.95 or greater indicate good model fit (Brown, 2006). After establishing model fit, differences in parameters of interest were tested by fixing parameters to zero and comparing model fit to a model in which parameters were freely estimated using a likelihood ratio difference test (Curran et al., 2010; Eid et al., 2015).

### Physiological data recording and preprocessing

#### Pupil dilation

We recorded participant’s pupil dilation as a proxy for central LC-NE activity (Reimer et al., 2014; Costa and Rudebeck, 2016; Joshi et al., 2016; Breton-Provencher and Sur, 2019; Deitcher et al., 2019; Zerbi et al., 2019) along with gaze position using an infrared video-based eye tracker (EyeLink 1000 desktop mount; monocular setup; SR Research Ltd.; Ottawa, Canada) with a spatial resolution of up to 0.25° and a sampling rate of 1000 Hz. A forehead- and chin rest 53.5 cm from the computer screen was used to minimize participants’ head movements during measurements. Participants were instructed to maintain central fixation throughout all experiments and compliance with this instruction was enforced at the beginning of every trial. Each experiment started with a (re)calibration of the eye tracking system using a standard 5-point grid. During (re)calibration, fixation errors were kept < 0.5°.

For synchronous, integrated analysis of eye tracking and EEG data we used the Eye-EEG toolbox (<u>Eye-EEG</u>; Dimigen, Sommer, Hohlfeld, Jacobs, & Kliegl, 2011), an extension for the open-source Matlab toolbox EEGLab (<u>EEGLAB</u>, RRID:SCR_007292; Delorme & Makeig, 2004) as well as the FieldTrip toolbox (<u>FieldTrip</u>, RRID:SCR_004849; Oostenveld, Fries, Maris, & Schoffelen, 2011). Eye tracking data of the re/conditioning and dichotic listening sessions was resampled to 500 Hz and segmented in bins of 8.5 s (i.e.,– 1.5–7 s with respect to CS onset). Time segments contaminated by blinks or excessive eye movements were automatically detected and imputed using custom-written Matlab code. In particular, segments falling more than three standard deviations below a participant’s median pupil dilation (calculated across the whole experiment) were considered as blink or partly occluded pupil. Further, periods with excessive eye movements as indicated by z-scored vertical gaze channel values > 3 (computed across the whole experiment) were considered as artifacts. All artifacts were padded by ± 50 samples to account for biased pupil estimates shortly before/after artifacts (cf. de Gee, Knapen, & Donner, 2014).

Excluding detected artifacts, we computed the average event-related pupil response for each trial category (i.e., CS+; CS–; separately for fear re/conditioning and dichotic listening data; see Figure 1c). In all trials, artifact containing segments were then replaced by the corresponding time segments of the demeaned average response centered at the given trial. Due to technical issues, no pupil data was available for one younger adult for fear conditioning on Day 2 and one older adult for fear conditioning on Day 3 and the dichotic listening task (Day 2).

Notably, we performed a set of control analyses that included a linear interpolation of missing pupil samples instead of the mean imputation approach described above. Importantly, irrespective of the preprocessing pipeline qualitatively similar results are obtained on the group level (see https://doi.org/10.17605/OSF.IO/G9FQJ).

#### Electroencephalography

To evaluate neural responses during re/conditioning and dichotic listening (cf. Figure 1c), we recorded the EEG. Data was continuously sampled from 61 Ag/AgCl electrodes embedded in an elastic cap that were placed according to the 10-10 system using BrainVision Recorder (BrainAmp DC amplifiers, Brain Products GmbH, Gilching, Germany; Braincap, BrainVision, respectively). An electrode above the forehead (AFz) served as ground. Three additional electrodes were placed next to each eye and below the left eye to acquire horizontal and vertical electrooculograms. Data was sampled at 1000 Hz in a bandwidth between 0.1–250 Hz and online-referenced to the right mastoid while the left mastoid was recorded as additional channel. During EEG-preparation, electrode impedances were kept <5 kΩ.

EEG data processing was performed by means of the Eye-EEG (Dimigen et al., 2011), EEGLab (Delorme and Makeig, 2004) and FieldTrip (Oostenveld et al., 2011) toolboxes in addition to custom-written Matlab code. For analyses, data was demeaned, re-referenced to mathematically linked mastoids, down-sampled to 500 Hz and band-pass filtered (0.2–125 Hz; fourth order Butterworth). A multi-step procedure was applied to purge data of artifacts: First, data was visually screened for periods of excessive muscle activity and subsequently independent component analysis (ICA) was used to identify and remove components related to eye, muscle and cardiac activity (e.g., Jung et al., 2000). Next, data was segmented in 8.5 s epochs (– 1.5 s and + 7 s with respect to stimulus onset) and submitted to a fully automatic thresholding approach for artifact rejection (cf. Nolan, Whelan, & Reilly, 2010). Excluded channels were interpolated with spherical splines (Perrin et al., 1989). Finally, remaining trials were again visually screened to determine successful cleaning.

Time-varying power information for each trial and electrode was then extracted by convolution of the cleaned time domain signal with a series of Morlet wavelets with a length of seven cycles (cf. Herrmann, Grigutsch, & Busch, 2005; Werkle-Bergner, Shing, Müller, Li, & Lindenberger, 2009). Time-varying power estimates were computed for frequencies between 1–40 Hz (in steps of 1 Hz) in a time window between –1.5 s to 7 s with respect to stimulus onset (time bins of 4 ms), separately for CS+ and CS– trials of the reconditioning and dichotic listening phases (see Figure 1c).

### Physiological data analyses

#### Within modality within-subject statistics (first level)

Within younger and older subjects, we contrasted arousing (CS+) and neutral control trials (CS–) by means of independent-samples *t*-tests to isolate arousal-associated response patterns. Contrasts were computed for time domain pupil data (i.e., Event-Related Pupil Response; ERPR), time domain EEG data (i.e., Event-Related Potential; ERP) and time-frequency domain EEG data (i.e., Event-Related Desynchronization; ERD). To counteract potential unequal distribution of CS+ and CS– trials (e.g., more artifacts in arousing trials), we iteratively selected random, equally sized subsets of the available trials using a bootstrapping procedure (n_bootstraps_ = 50 iterations; n_SelectedTrials_ = lowest trial number across conditions – 1). The mean *t*-value over the 50 bootstraps served as final first level test statistic that was passed on to the second level (see below). First level statistics were computed within subjects for conditioning (separately for each day (Day 1–3); see Figure 1c), reconditioning and dichotic listening trials. While the contrast (CS+ vs. CS–) remained the same across these analyses, please note that during re/conditioning participants received electrical stimulation (US) and thus the observed responses may represent a mixture of fear and somatosensory responses. In contrast, during the dichotic listening task no shocks were applied and thus observed responses (CS+ vs CSin–) indicate the reinstatement of the fear response (cf. Figure 1c).

Notably, the CS+ vs. CS– first level *t*-maps express the difference in pupil and EEG responses to arousing and non-arousing stimuli. That is, in the strict sense of the word they do not constitute an ERPR / ERP /ERD, but rather the standardized difference between two pupil / EEG responses. However, to ease readability, we use the terms ERPR, ERP and ERD also for these contrasts.

#### Within-modality group statistics (second level)

For analyses on the group level, we contrasted first level *t*-maps (i.e., CS+ vs CS–) against zero to identify neural correlates associated with the arousal manipulation in conditioning and reconditioning trials (cf. Figure 1c) that were shared across subjects in each group. Analyses were run separately for each day, first across all subjects (YA and OA) and then within YA and OA for all modalities (ERPR, ERP and ERD data). In particular, we calculated non-parametric, cluster-based, random permutation tests as implemented in the FieldTrip toolbox that effectively control the false alarm rate in case of multiple testing (Maris and Oostenveld, 2007; Oostenveld et al., 2011). Please note that the same statistical procedure was applied to two-dimensional (i.e., channel × time) and three-dimensional (i.e., channel × frequency × time) data. That is, ERPR, ERP and ERD were analyzed in the same manner. Here, however, only the approach for three-dimensional data (ERD) is described to ease readability. In short, first a two-sided, dependent samples *t*-test was calculated for each spatio-spectral-temporal (channel × frequency × time) sample. Neighboring samples with a *p*-value below 0.05 were grouped with spatially, spectrally and temporally adjacent samples to form a cluster. The sum of all *t*-values within a cluster formed the respective test-statistic (*t*_sum_). A reference distribution for the summed cluster-level *t*-values was computed via the Monte Carlo method. Specifically, in each of 1000 repetitions, group membership was randomly assigned, a *t*-test computed and the *t*-value summed for each cluster. Observed clusters whose test-statistic exceeded the 97.5^th^ percentile for its respective reference probability distribution were considered significant.

On a group level, cluster statistics revealed reliable arousal effects during conditioning and reconditioning within all modalities (ERPR, ERP, and ERD). To evaluate a potential reinstatement of these fear responses also during the dichotic listening task (in which no shocks were applied anymore; cf. Figure 1c), each subjects’ first level (CS+ vs. CS–) dichotic listening data was averaged across spatio-spectral-temporal samples that reached significance on a group level during the reconditioning period. That is, we applied the observed reconditioning fear response (i.e., significant cluster) as a search space to evaluate its reinstatement within the dichotic listening data. This approach yielded a single reinstatement value for each subject for each modality (i.e., ERPR, ERP and ERD data). Within modalities, the reliability of the reinstatement was then determined by means of non-parametric Wilcoxon signed rank (W) tests (across and within age groups).

To judge the temporal stability of fear conditioned pupil responses (ERPR) over assessment days (Day 1–3; cf. Figure 1c), in addition each subjects’ first level (CS+ vs. CS–) conditioning and reconditioning ERPR data was averaged across time points that reached significance on a group level (i.e., second level statistic for the respective day). This yielded a single ERPR value for each subject for each conditioning session (Day 1–3) and the reconditioning phase (Day 2). We then used intra-class-correlations (ICC; two-way mixed; consistency) to evaluate the temporal stability of participants’ fear conditioned pupil dilation.

#### Cross-modality group statistics

To determine whether EEG correlates of the arousal manipulation (i.e., ERP, ERD) were linked to the LC-NE system, we correlated participants’ EEG responses with their pupil dilation, a proxy of noradrenergic activity, across age groups. We assessed this association within the dichotic listening data, since this provides an estimate of the reinstatement of the fear response irrespective of potential somatosensory artifacts related to the reinforcement (US; see Figure 1). In particular, participants’ first level EEG *t*-maps (CS+ vs. CS–) were correlated with participants’ average pupil reinstatement (see within modality group statistics) in a non-parametric, cluster-based, random permutation framework as implemented in the FieldTrip toolbox. Analyses were run separately for ERP and ERD data, however, here only the approach for time-frequency data (ERD) is described to ease readability. For each spatio-spectral-temporal sample a two-sided Pearson’s correlation between the EEG and the pupil reinstatement data was calculated. As done for the within-modality statistics, neighboring samples with a *p*-value below 0.05 were grouped with spatially, spectrally and temporally adjacent samples to form a cluster. The sum of all *rho*-values within a cluster formed the respective test-statistic. A reference distribution for the summed cluster-level *rho*-values was computed via the Monte Carlo method. In particular, the null hypothesis of statistical independence between EEG and pupil data was tested by randomly permuting pupil estimates between subjects over 1000 repetitions. For each repetition, a correlation was computed and *rho*-values were summed for each cluster. Observed clusters whose test-statistic exceeded the 97.5^th^ percentile for its respective reference probability distribution were considered significant. To specifically target reinstatement responses, we restricted the cross-modality analyses to EEG samples that showed a reliable arousal effect (i.e., were part of significant clusters in the second level reconditioning analyses). Note however, that analyses were performed solely on reinstatement data (i.e., dichotic listening ERPR, ERP and ERD data; cf. Figure 1).

#### Cross-modality structural equation model

Cluster-correlation analyses revealed reliable associations between EEG and pupil reinstatement for both ERP and ERD data, suggesting a common underlying pupil– EEG factor. For each subject, we thus extracted and averaged those samples of the EEG reinstatement response that showed a reliable link to pupil reinstatement. This returned a single pupil-associated reinstatement estimate for ERP and ERD data, respectively. As for the behavioral data, we then used a structural equation modeling approach to integrate over the interrelated indicators of the arousal response (see Figure 2, upper part). In particular, in younger and older adults, standardized pupil reinstatement and pupil-associated EEG reinstatement estimates served as manifest variables and loaded on a single, latent NE responsiveness factor (i.e., a multiple-group model). Factor loadings (other than the first, which was fixed to one) were estimated freely but were constrained to be equal across groups. The model demonstrated strict factorial invariance (i.e., showed invariant manifest intercepts and variances across groups) and thus allowed an interpretation of age group differences in the means of the latent factor (Meredith and Teresi, 2006; Schwab and Helm, 2015). We evaluated age differences in latent NE responsiveness by means of Spearman correlations (across age groups). Adequacy of the proposed model was assessed using a *χ*^2^-test as well as two additional fit indices (RMSEA, CFI; see above).

#### Analyses of associations between physiological and behavioral data

After generating structural equation models for our cognitive and physiological measures, respectively, we set out to link both modalities. That is, we were interested in assessing the relation between interindividual differences in attention and interindividual differences in NE responsiveness. For this, we first built a unified model merging the attention and responsiveness models described above (see Figure 2). We then investigated associations between cognitive and physiological factors by allowing for freely estimated covariances on a latent level (shown in yellow, Figure 2). As before, model fit for all described models was determined using a *χ*^2^-test in combination with two additional fit indices (RMSEA, CFI).

### Code and data availability

The custom code and (anonymized) data used for these analyses is available from the corresponding authors upon request.

## Results

### Impaired selective attention in older adults

Participants demonstrated successful auditory selective attention in the dichotic listening task as indicated by a two-factorial mixed measures ANOVA (Age group × Attentional Focus; main effect of attentional focus: *F*(1, 75) = 26.413, *p <* 0.001, *η*² = 0.260). Post-hoc analyses within younger and older adults demonstrated that both groups were able to exert auditory selective attention (one-factorial repeated measures ANOVA; main effect of attentional focus: For YA *F*(1, 38) = 22.803, *p <* 0.001, *η*² = 0.375; For OA *F*(1, 37) = 5.702, *p* = 0.022, *η*² = 0.134).

Younger and older adults, however, differed reliably in their ability to modulate their attentional focus. While the Age group main effect was not significant (*F*(1, 75) = 0.087, *p =* 0.769, *η*² = 0.001); we observed a reliable Age group × Attentional focus interaction (*F*(1, 75) = 16.318, *p <* 0.001, *η*² = 0.179; see Figure 3) indicating impaired selective attention in old age. Please note, that here a main effect of Age (e.g., lower LI in older compared to younger adults) would indicate better per performance in one Attentional focus condition (e.g., FL) and worse performance in the other (e.g., FR; cf. Figure 3). The observed Age group × Attentional focus interaction in contrast reveals worse performance in older adults in both conditions (i.e., lower LI values in the FR condition; higher LI values in the FL condition).

**Figure 3.**
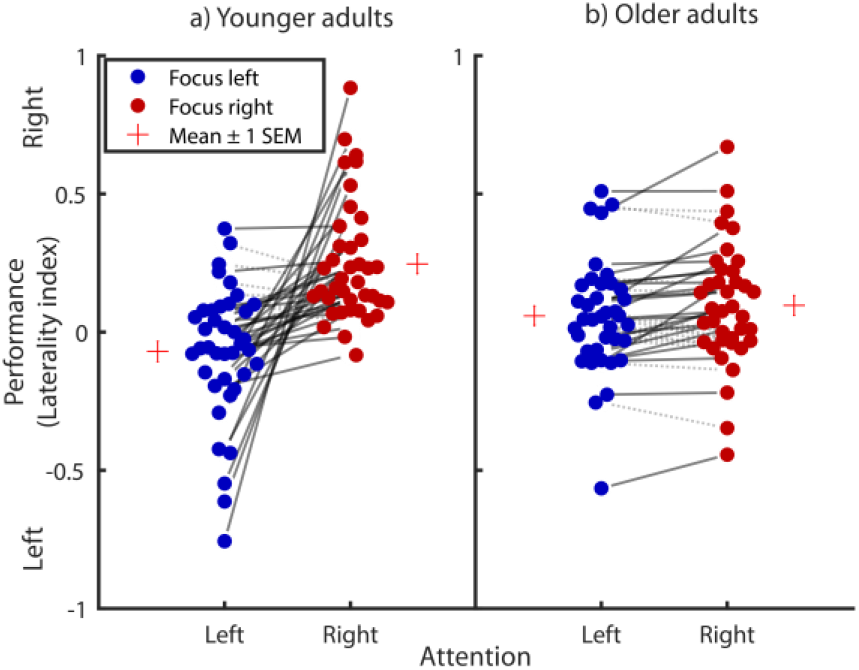
Selective auditory attention performance of younger (a) and older adults (b) in a dichotic listening task. Negative and positive laterality index values indicate a tendency for left and right ear responses, respectively. Circles connected by solid lines show participants who demonstrate a behavioral selective attention effect (i.e., more responses of the cued ear relative to the not cued ear), with the slope of the lines reflecting the degrees of attentional modulation. Circles connected by grey dotted lines indicate a reversed effect. While the amount of selective attention is markedly decreased in older adults, both younger and older participants demonstrate reliable selective attention on a group level. SD = Standard Deviation; Laterality index = (Focus Right – Focus Left) / (Focus Right + Focus Left)).

Post-hoc analyses indicated that age differences in auditory selective attention were not explained by age related differences in hearing loss (i.e., Age group × Attentional Focus mixed measures ANOVA including hearing loss as covariate: Age group × Attentional focus: *F*(1, 74) = 4.862, *p =* 0.031, *η*² = 0.062; Hearing loss × Attentional focus: *F*(1, 74) = 0.740, *p =* 0.393, *η*² = 0.010). Similarly, control analyses indicated that age differences in auditory selective attention persisted after including interaural threshold differences in the model (see: https://doi.org/10.17605/OSF.IO/G9FQJ).

We replicated this finding of impaired selective attention in aging across multiple visual and auditory attention tasks using non-parametric Mann-Whitney U-tests (all *p*s < 0.05; see Table 2). In order to later reliably relate attention performance to physiological indices of the LC-NE system (see below), we integrated performance over tasks to derive a single measure reflecting general attention performance (see Figure 2, lower part). The proposed model fit the data well (*χ*^2^ = 9.827, *df* = 23; RMSEA = 0.0; CFI = 1.594; Brown, 2006). The variances of the attention factors differed reliably from zero in both age group (all Δ*χ*^2^ ≥ 11.225, Δ*df* = 1, all *p* < 0.001) indicating interindividual differences in attention.

### Stable fear conditioned pupil dilation in younger and older adults

In the conditioning and reconditioning phases of the assessment (see Figure 1c), younger and older adults demonstrated a reliable, multimodal response to the arousal manipulation. In the following, first modality-specific results are reported (i.e., pupil dilation and EEG) before detailing their interrelation.

During fear conditioning and reconditioning, conditioned stimuli (CS+ vs. CS–) reliably elicited pupil dilation over prolonged time windows as revealed by cluster permutation analyses (both across and within age groups all *p*s_corr_ < 0.01; see Figure 4 and also https://doi.org/10.17605/OSF.IO/G9FQJ).

**Figure 4.**
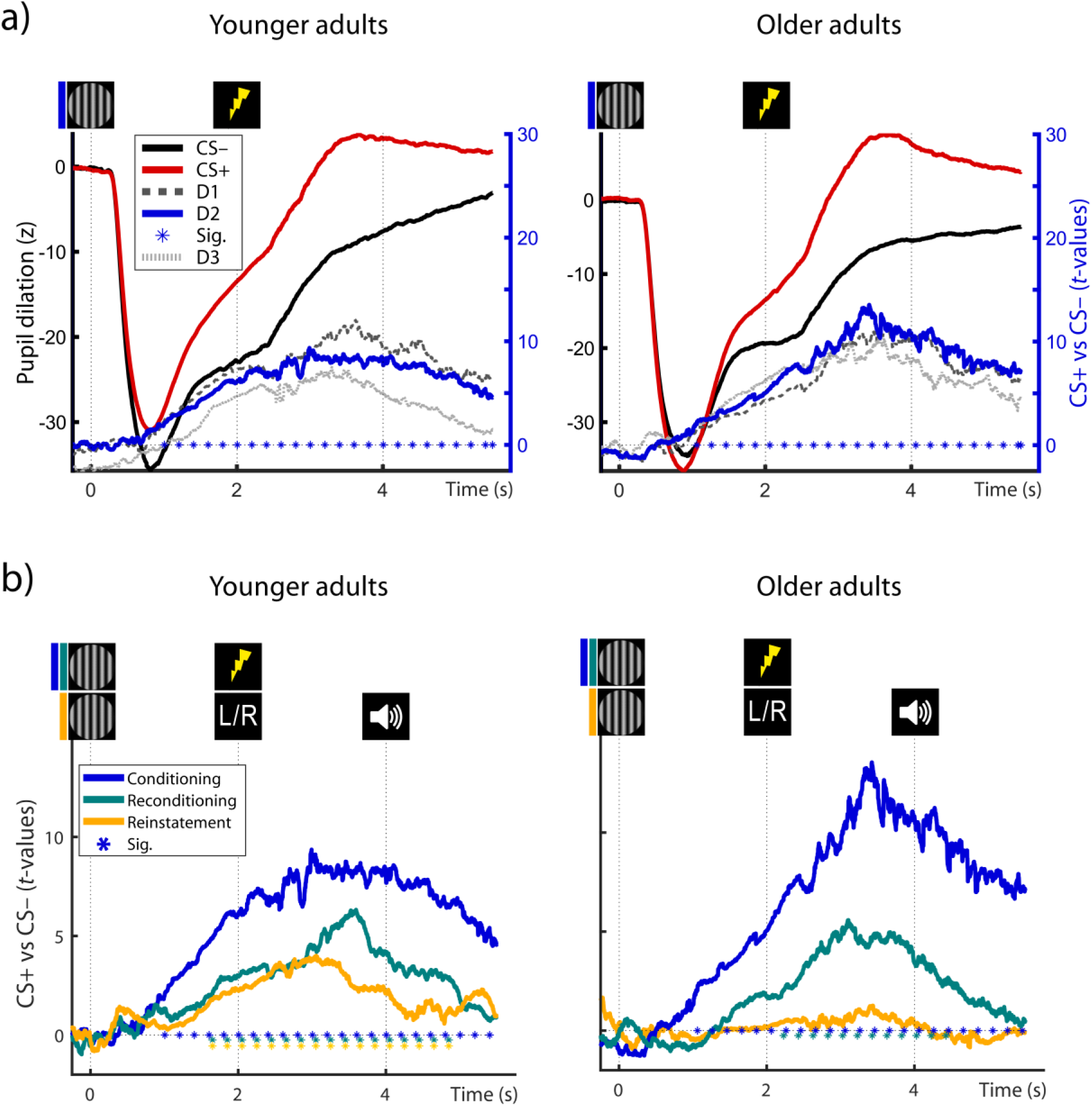
Average pupil dilation of younger and older adults in response to the presentation of fear conditioned (CS+; red) and neutral control stimuli (CS–; black) during fear conditioning (a; Day 2) and during the dichotic listening task (b; i.e., reconditioning and reinstatement). Group statistics depict the consistency of the CS+ vs CS– contrast on the second level. **a.** Statistics are presented for conditioning data for the first (D1; light grey, dashed), second (D2; solid, blue) and third (D3; dark grey, dotted) day of conditioning assessments (see right y-axis). **b**. Pupil responses during conditioning (blue), reconditioning (teal) and dichotic listening trials (reinstatement; orange) on day 2. Horizontal lines of asterisks indicate significant time windows (blue *, Sig.). Reinstatement of the fear conditioned pupil response during the dichotic listening task (see Figure 1c) is evaluated statistically using Wilcoxon tests (YA: *W*(39) = 586; *Z* = 2.735; *p* = 0.006; OA: *W*(37) = 364; *Z* = 0.189; *p* = 0.850).

Fear conditioned pupil responses demonstrated a moderate to high stability across assessments (Day 1–3) as indicated by intra-class-correlations (two-way mixed; consistency; ICC (95%CI) = 0.652 (0.502–0.766); p < 0.001). In line with pupil dilation as a non-invasive marker of LC activity, this points to a stable phasic activation of the NE system by fear conditioned stimuli (and US) across age groups.

### Reinstatement of pupil dilation in younger adults

In the absence of reinforcements (US), fear conditioned stimuli maintained their arousing nature and led to a marginally significant reinstatement of pupil dilation across groups (*W*(76) = 1839; *Z* = 1.948; *p* = 0.052). While younger adults demonstrated a robust reinstatement effect, in older adults reinstatement did not reach statistical significance on a group level (see Figure 4; YA: *W*(39) = 586; *Z* = 2.735; *p* = 0.006; OA: *W*(37) = 364; *Z* = 0.189; *p* = 0.850). The lack of pupil reinstatement in older adults presumably reflects age-related difficulties in triggering and maintaining self-initiated processing (i.e., reinstatement; Lindenberger and Mayr, 2014) in line with previous reports (van Gerven et al., 2004). By contrast, age differences are known to be reduced or even disappear when older adults can rely on external information (e.g., reminders), like the reinforcements (US) during re/conditioning. The age difference in the reinstatement of pupil dilation approached statistical significance (YA vs. OA: *U*(76) = 1671; *Z* = 1.756; *p* = 0.079). Since the reinstatement of pupil dilation occurs in the absence of somatosensory stimulation and associated artifacts, it is attributed to the arousal response following the reactivation of the fear memory. We thus interpret the reinstatement of the fear-induced pupil response as indicator of the effectiveness of the LC-NE system in modulating memory, which trends to be reduced in aging.

### Fear conditioned parietal event-related potentials in younger and older adults

During fear reconditioning, conditioned stimuli also reliably elicited event-related EEG responses (ERP) both across and within age groups as revealed by cluster permutation analyses (all *p*s_corr_ < 0.01; see Figure 5 and https://doi.org/10.17605/OSF.IO/G9FQJ). In particular, we observed that after an initially similar ERP (< 1 s) to CS+ and CS–, conditioned stimuli (CS+) were associated with an increasingly more negative going slow wave in the delay interval (between CS+ (t = 0 s) and US onset (t=2 s)). This was reflected in a sustained negative cluster with strongest polarity at centro-parietal electrodes (i.e., parietal slow wave; see Figure 5). Following the onset of the reinforcement (US), in CS+ trials the ERP rapidly flipped its polarity while maintaining a highly similar parietal topography, thus giving rise to a sustained positive cluster (i.e., late parietal potential; see Figure 5). In line with the established role of anticipatory slow waves and late parietal potentials in arousal and emotion processing (for reviews see Schupp, Flaisch, Stockburger, & Junghöfer, 2006 and van Boxtel & Böcker, 2004; Vazey et al., 2018), this points to increased sustained attention to CS+ during the anticipatory delay interval (0–2 s) and an augmented arousal response following US (> t = 2 s). Both the topography and time course of the ERP responses were highly similar across age groups, indicating a maintained arousal response to conditioned stimuli (CS+; and US) across the lifespan.

**Figure 5.**
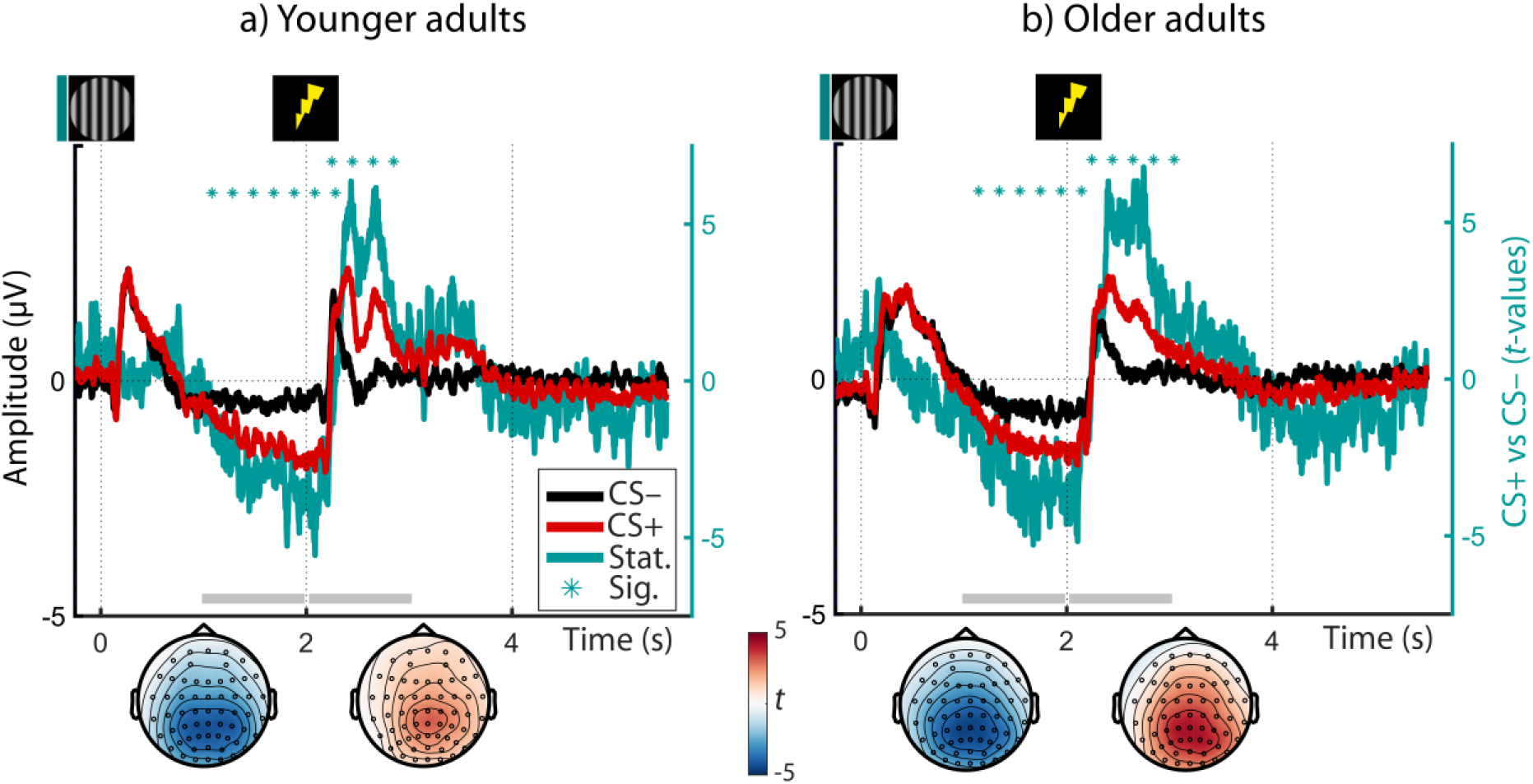
Averaged Event-Related Potentials (ERP) of younger (a) and older adults (b) in response to the presentation of fear conditioned (CS+; red) and neutral control stimuli (CS–; black) during reconditioning (left y-axis; on Day 2). Group statistics depict the consistency of the CS+ vs CS– contrast on the second level (teal) and are shown on the right y-axis. Horizontal lines of asterisks indicate significant time windows (*, Sig.). The topography of the group statistics between 1–2 and 2–3 s relative to CS onset is shown below the time courses (gray horizontal bars). For visualization, time courses are averaged across all electrodes.

Notably, as the earlier anticipatory potential emerges before onset of the reinforcement, we can rule out that it constitutes an artifact of the somatosensory stimulation. The latter potential in contrast may be influenced by the reinforcement and should only be considered further if it is reliably reinstated in absence of the stimulation (see below).

### Reinstatement of parietal event-related potentials in younger and older adults

ERP to the arousal manipulation were reinstated in the dichotic listening task in the absence of reinforcements (US). Across younger and older adults, both the earlier, anticipatory negative potential as well as the later, positive potential reached significance (Negative: *W*(77) = 861; *Z* = –3.252; *p* = 0.001; Positive: *W*(77) = 2028; *Z* = 2.673; *p* = 0.008). While in younger adults, only the anticipatory response was reliably reinstated (Negative: *W*(39) = 214; *Z* = –2.456; *p* = 0.014; Positive: *W*(39) = 456; *Z* = 0.921; *p* = 0.357; see Figure 6), in older adults both reached statistical significance (Negative: *W*(38) = 214; *Z* = –2.270; *p* = 0.023; Positive: *W*(38) = 582; *Z* = 3.067; *p* = 0.002; see Figure 6). The age difference in the latter, positive response was marginally significant (YA vs. OA: *U*(77) = 1335; *Z* = –1.890; *p* = 0.059). In the absence of somatosensory stimulation and associated artifacts, the reinstatement of the parietal reconditioning ERPs is attributed to the arousal response following the reactivation of the fear memory (van Boxtel and Böcker, 2004; Schupp et al., 2006).

**Figure 6.**
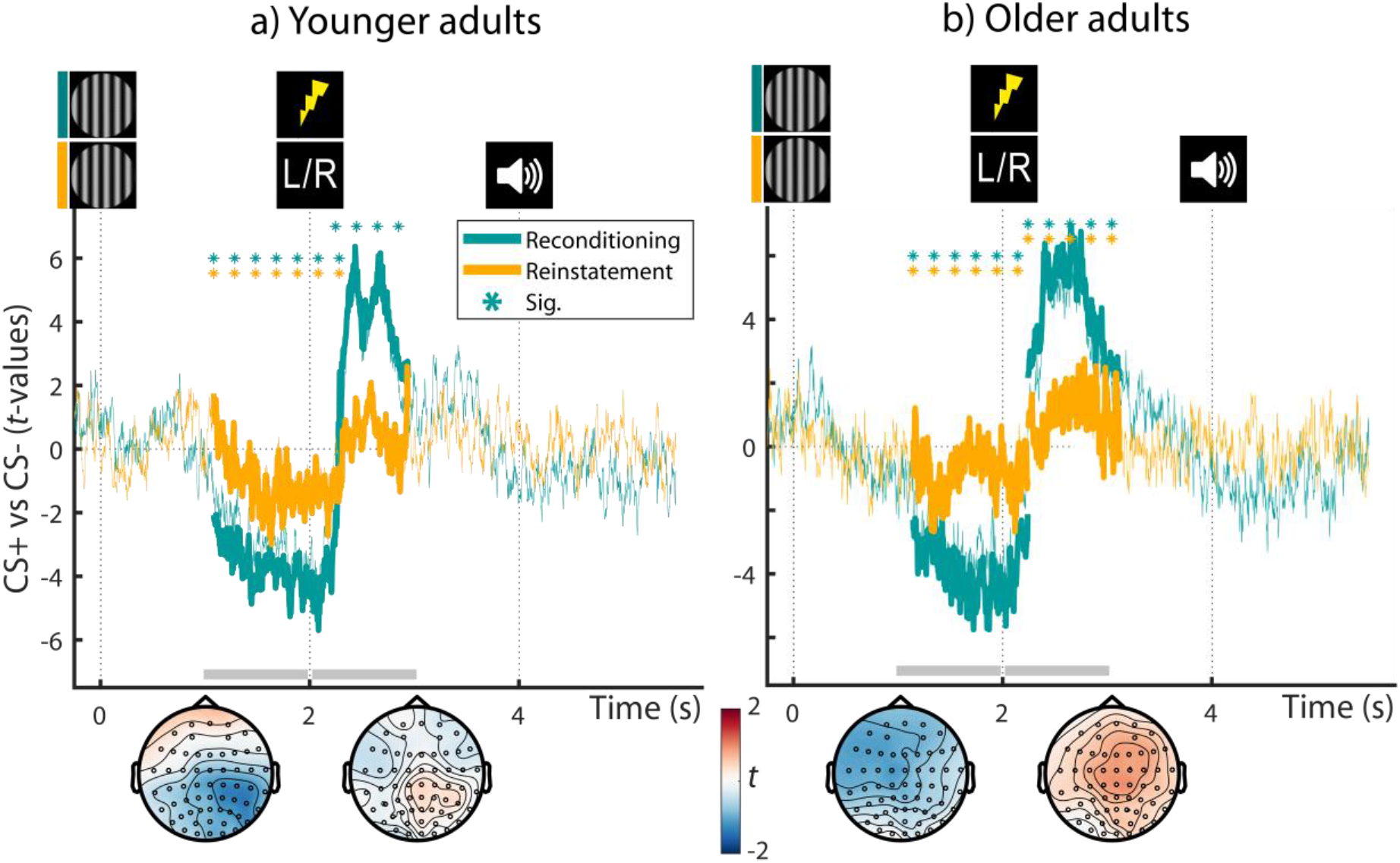
Group statistics for younger (a; YA) and older adults’ (b; OA) Event-Related Potentials (ERP) during reconditioning (teal; cf. Figure 5), and dichotic listening trials (orange; reinstatement) on day 2. All statistics depict the consistency of the CS+ vs CS– contrast on the second level. Teal horizontal lines of asterisks (*; Sig.) indicate the extent of the significant reconditioning clusters. Reinstatement of the earlier, negative and later, positive ERP clusters during the dichotic listening task is evaluated statistically using Wilcoxon tests (see orange lines of asterisks; YA: Negative: *W*(39) = 214; *Z* = –2.456; *p* = 0.014; Positive: *W(*39) = 456; *Z* =0.921; *p* = 0.357; OA: Negative: *W*(38) = 214; *Z* = –2.270; *p* = 0.023; Positive: *W*(38) = 582; *Z* =3.067; *p* = 0.002). The topography of the reinstatement group statistics between 1–2 and 2–3 s relative to CS onset is shown below the time courses (gray horizontal bars). CS+ = fear conditioned stimulus; CS– = perceptually matched, neutral control stimulus. For visualization, time courses are averaged across all electrodes.

### Fear conditioned posterior desynchronization in younger and older adults

Across age groups, fear conditioned stimuli were also associated with a sustained (t = –0.248–4.956 s) decrease in low EEG frequencies (4–40 Hz). The observed cluster was most pronounced in the theta to beta frequency bands (∼5–25 Hz) and showed its strongest polarity at parieto-occipital electrodes (cf. van Boxtel & Böcker, 2004; YA and OA: *p*_corr_ = 0.002). Analyses within younger and older adults revealed reliable desynchronization effects in both age groups with strongest extent at posterior electrodes (YA: *p*_corr_ = 0.024; OA: *p*_corr_ = 0.002; see Figure 7). However, in younger adults desynchronization was restricted to the anticipatory delay phase (i.e., before US onset; t = 0.352–1.360 s; 5–40 Hz) whereas in older adults a more persistent desynchronization was observed (t = –0.248–5.144 s; 2–40 Hz). In addition, coinciding with onset of the reinforcement (US), a positive low frequency cluster emerged across and within age groups (see Figure 7). However, this positive cluster most likely reflects an artifact of the electric stimulation (2 Hz pulse at T = 2 s) and consequently was not evident in absence of the US (see Figure 8, below).

**Figure 7.**
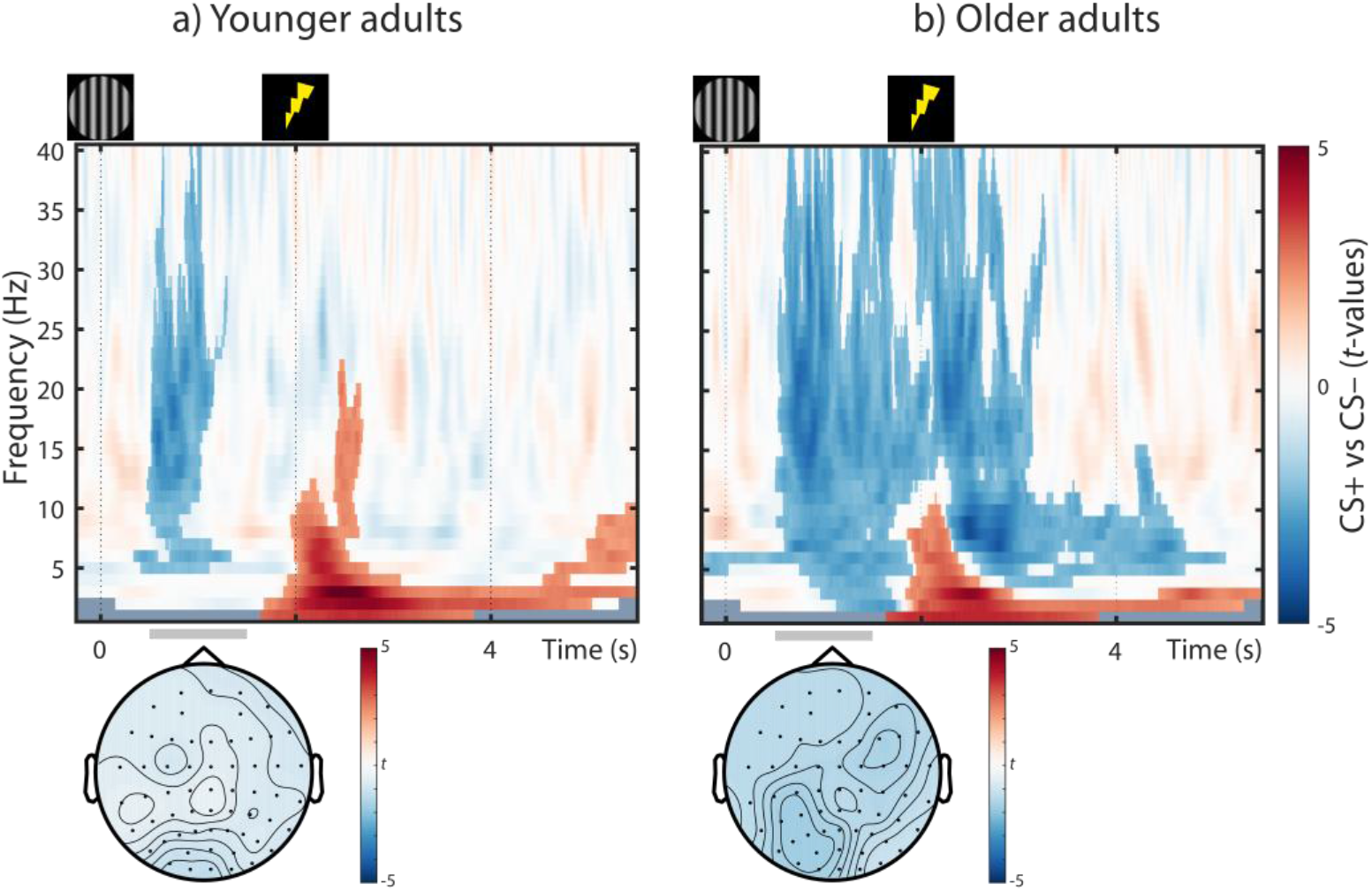
Group statistics for younger (a) and older adults’ (b) time-frequency EEG responses during reconditioning trials on day 2 (i.e., Event-Related Desynchronization). Statistics depict the consistency of the CS+ vs. CS– contrast on the second level. Non-significant samples are displayed with 50% transparency, while significant clusters are overlaid without transparency. The topography between 0.5–1.5 s relative to CS onset is shown below the time courses (gray horizontal bars). For topographies, data is averaged across all frequencies.

**Figure 8.**
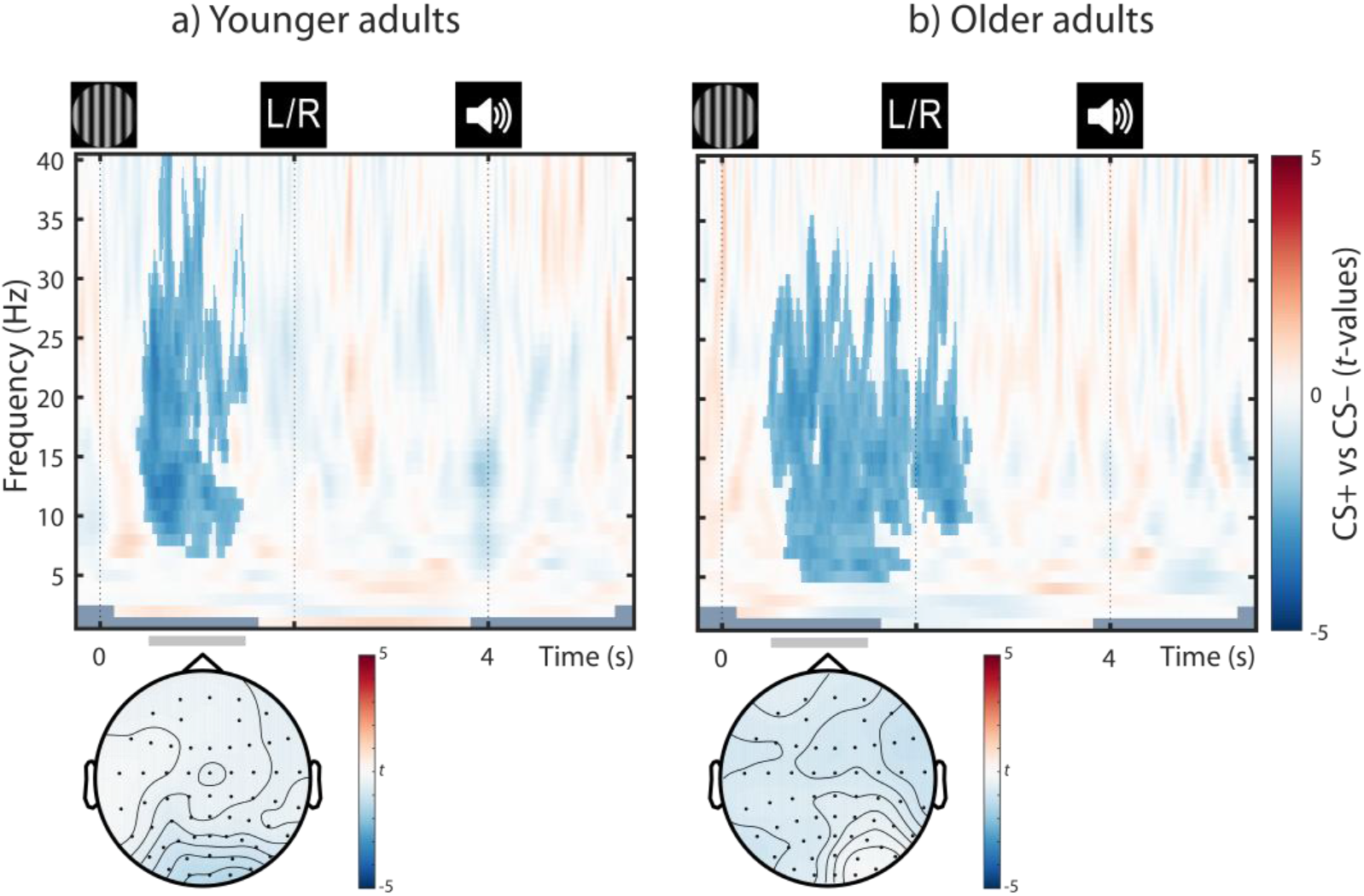
Group statistics for younger (a) and older adults’ (b) time-frequency EEG responses during dichotic listening trials on day 2 (i.e., reinstatement of the Event-Related Desynchronization). Statistics depict the consistency of the CS+ vs CS– contrast on the second level. For visualization purposes, non-significant samples are displayed with 50% transparency, while significant clusters are overlaid without transparency (YA: *p*_corr_ = 0.044; OA: *p*_corr_ = 0.004). The topography between 0.5–1.5 s relative to CS onset is shown below the time courses (gray horizontal bars). For topographies, data is averaged across all frequencies.

Desynchronization in low frequencies has been associated with NE-associated changes in cortical and behavioral state (McCormick et al., 1991; Harris and Thiele, 2011; Marzo et al., 2014; Safaai et al., 2015; Neves et al., 2018). In particular, in humans desynchronization in the alpha–beta band was linked to facilitated information processing resulting from a decrease in cortical inhibition (for reviews, see Hanslmayr, Staudigl, & Fellner, 2012; Jensen & Mazaheri, 2010; Klimesch, Sauseng, & Hanslmayr, 2007). Accordingly, the pronounced low frequency desynchronization in response to conditioned stimuli (CS+; and US) suggests an anticipatory transition towards a more activated cortical state, including increased cortical excitability and attention deployment. For spectral responses to the arousal manipulation split by frequency band, please see : https://doi.org/10.17605/OSF.IO/G9FQJ

### Reinstatement of posterior desynchronization in younger and older adults

Presentation of fear conditioned stimuli (CS+) during the dichotic listening task (i.e., without reinforcements; US) reinstated a pronounced decrease in low frequencies across age groups (YA and OA: *W*(77) = 900; *Z* = –3.054; *p* = 0.002). Similarly, within younger adults we observed a reliable reinstatement with a mostly posterior topography (YA: *W*(39) = 161; *Z* = –3.196; *p* = 0.001), whereas older adults showed a marginally significant reinstatement with a more wide-spread extent (OA: *W*(38) = 239; *Z* = –1.907; *p* = 0.057; see Figure 8). As for the pupillary and parietal ERP reinstatement, the reinstatement of the low frequency desynchronization is considered as arousal response to the reactivated fear memory.

### Reinstatement of EEG arousal responses is associated with pupil dilation

To briefly summarize the modality-specific findings, across groups we observed a negative, anticipatory slow wave and a late, positive parietal potential (ERP; see Figure 5; van Boxtel and Böcker, 2004; Schupp et al., 2006), a low frequency desynchronization (ERD; see Figure 7; McCormick et al., 1991; Harris and Thiele, 2011; Marzo et al., 2014), as well as pupil dilation (ERPR; see Figure 4; Joshi et al., 2016; Reimer et al., 2016; Breton-Provencher and Sur, 2019; Deitcher et al., 2019; Zerbi et al., 2019). Within the dichotic listening task (i.e., in the absence of reinforcements; US; see Figure 1) we largely witnessed a reliable reinstatement of the arousal response across modalities (i.e., ERP, ERD, ERPR; see Figures 4, 6, and 8). We interpret the reinstatement of the arousal response as reflecting a phasic activation of the LC-NE system by the reactivated fear memory. To support this claim, electrophysiological reinstatement marker (ERP, ERD) should be linked to pupil reinstatement, a non-invasive index of locus coeruleus activity.

Accordingly, cluster permutation correlations revealed a reliable association between the reinstatement of the anticipatory, parietal slow wave and pupillary reinstatement (*p*_corr_ = 0.028). The cluster reached significance in the time window previously filled by the reinforcement (during (re)conditioning; t = 2.108–2.212 s; see Figure 1 and 9) and showed its strongest polarity at left lateralized centro-parietal electrodes. Spearman correlation coefficients are reported in Table 3 for analyses across and within groups (for this, EEG data was averaged across the cluster). No reliable link was observed between the second, positive ERP cluster (late parietal potential) and pupil dilation (t > 2 s; *p*_corr_ > 0.1). This indicates that the late parietal potential was not reliably linked to our LC-NE activity index and we thus dropped it from further analyses.

**Figure 9.**
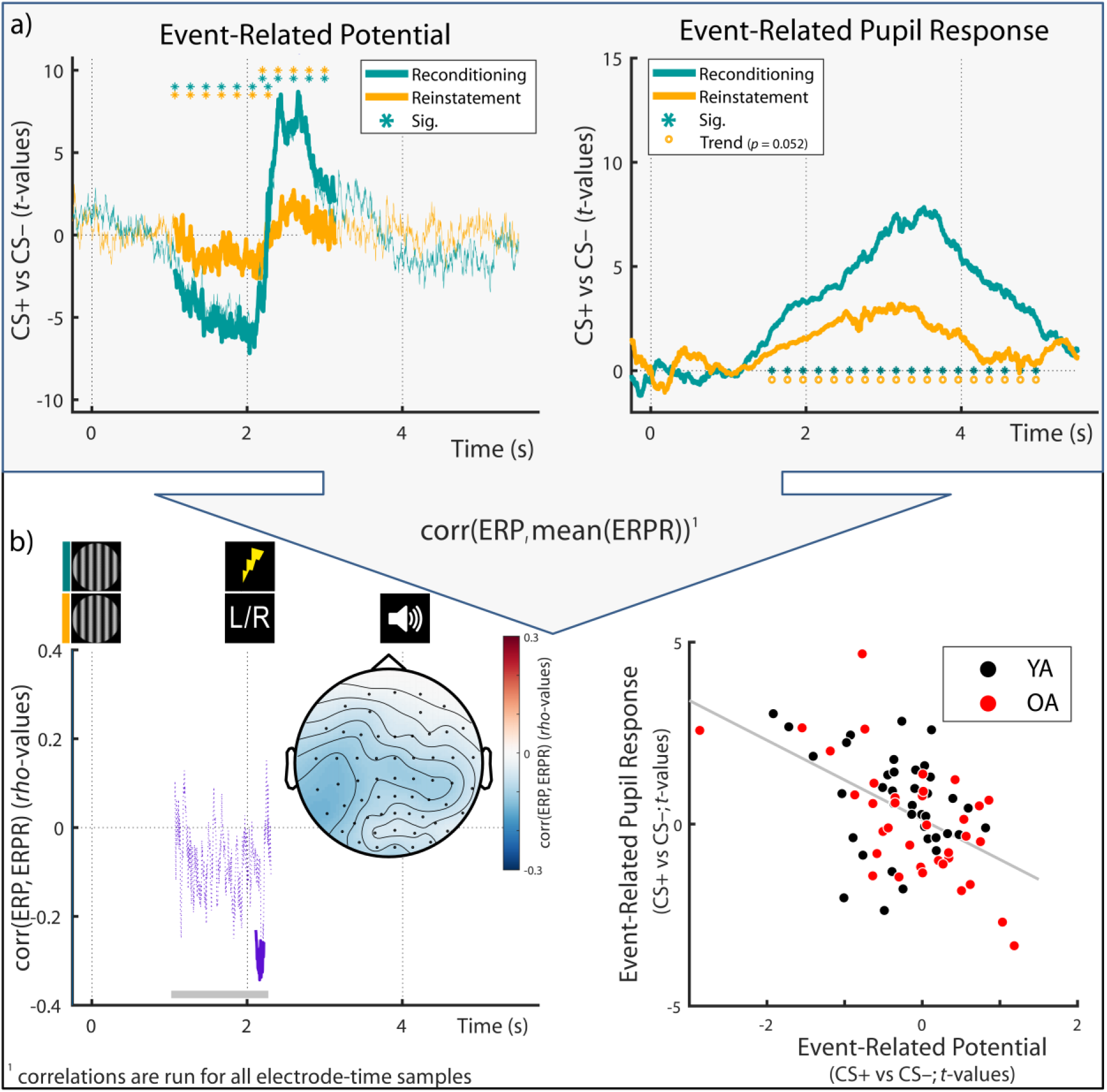
(a) Across age groups, fear conditioned Event-Related Potentials (ERP) and Event-Related Pupil Responses (ERPR) to the arousal manipulation learned during re/conditioning (teal lines) were reinstated in the dichotic listening task (organge lines in the left and right panel, respectively). For ERPR and ERP analyses within age-groups, see Fig. 4 and 5–6, respectively. (b; left panel) Within electrode and time ranges that demonstrated a reliable arousal effect during reconditioning, ERP and ERPR reinstatement data were correlated. A significant negative association was observed between t = 2.108–2.212 s (solid purple line). The topography between 1–2.3 s relative to CS onset (see gray horizontal bars) is shown below the time course. (b; right panel) For visualization purposes only, an additional scatter plot is provided depicting the same association between ERP and ERPR reinstatement data (here ERP data is averaged over those samples [time, electrodes] that formed the reliable cluster [see left panel for cluster extend]). Corr: Correlation; YA: Younger adults; OA: Older adults; CS+: Conditioned stimulus; CS–: Neutral control stimulus, perceptually matched to the CS+

**Table 3.** Overview of associations between pupillary and EEG reinstatement components

| Measure | Group | (N) | *rho* | *p* |
| --- | --- | --- | --- | --- |
| Parietal slow wave | YA + OA | (N = 76) | -0.424 | <0.001 |
| Alpha-beta desynchronization | YA + OA | (N = 76) | -0.424 | <0.001 |
| Parietal slow wave | YA | (N = 39) | -0.255 | 0.116 |
| Alpha-beta desynchronization | YA | (N = 39) | -0.401 | 0.011 |
| Parietal slow wave | OA | (N = 37) | -0.567 | <0.001 |
| Alpha-beta desynchronization | OA | (N = 37) | -0.411 | 0.011 |

Note: Pupil and EEG reinstatement was assessed in the dichotic listening task (cf. Figure 1c). The first row within age groups reports Spearman's correlations of pupil dilation and the anticipatory time domain EEG cluster while the second row reports the time-frequency domain EEG cluster. YA: Younger adults; OA: Older adults.

Moreover, a stronger reinstatement of low EEG frequency desynchronization in response to conditioned stimuli (CS+ vs. CS–) was associated with a larger reinstatement of pupil dilation (*p*_corr_ = 0.022; see Figure 5b). The cluster reached significance in the anticipatory delay phase (i.e., before US onset during (re)conditioning; t = 0.588–1.160 s) in the alpha–beta frequency band (9–30 Hz) and was most pronounced at parieto-occipital electrodes (see Figure 10). Spearman correlation coefficients (based on EEG data averaged across the cluster) are provided in Table 3.

**Figure 10.**
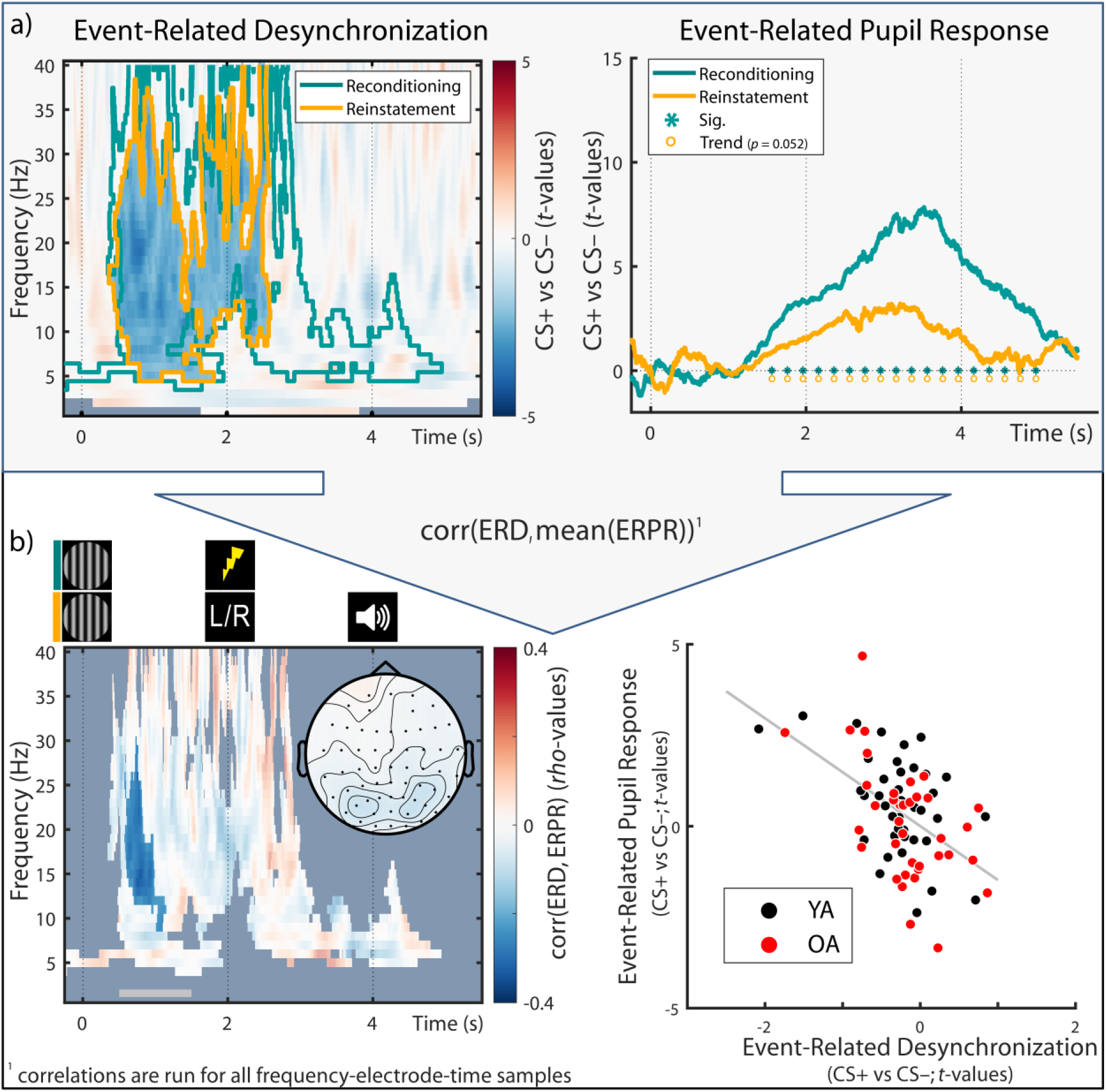
(a) Across age-groups, fear conditioned Event-Related Desynchronization (ERD) and Event-Related Pupil Responses (ERPR) to the arousal manipulation learned during re/conditioning (teal (out)lines) were reinstated in the dichotic listening task (orange (out)lines in the left and right panel, respectively). For ERPR and ERD analyses within age-groups, see Fig. 4 and 7–8, respectively. (b; left panel) Within electrode, time and frequency-ranges that demonstrated a reliable arousal effect during reconditioning, ERD and ERPR reinstatement data were correlated. A significant negative association was observed between t = 0.558–1.160 s (non-transparent cluster). The topography between 0.5–1.5 s relative to CS onset (see gray horizontal bar) is shown below the time course. (b; right panel) For visualization purposes only, an additional scatter plot is provided depicting the same association between ERD and ERPR reinstatement data (here ERD data is averaged over those samples [time, electrodes, frequencies] that formed the reliable cluster [see left panel for cluster extend]). Corr: Correlation; YA: Younger adults; OA: Older adults; CS+: Conditioned stimulus; CS–: Neutral control stimulus, perceptually matched to the CS+

In sum, in line with our interpretation cluster correlations indicated that both ERP and ERD responses to the arousal manipulation were linked to pupil dilation, a proxy for LC-NE activity. To further corroborate this conclusion, we repeated our analyses, this time using reconditioning instead of reinstatement pupil and EEG data (cf. Figure 1c). We again observed a reliable, qualitatively similar, association between EEG responses and pupil dilation, suggesting a common dependence on LC-NE activity (ERP: *p* = 0.026 and *p* = 0.044 (two reliable clusters), t = 1.12–1.218 s and t = 1.706–1.832 s; ERD: *p* = 0.03, t = 0.648–1.112 s, frequency range = 13–37 Hz). Crucially, we additionally repeated these analyses on the single-trial level. That is, we tested whether we could predict a given trial’s phasic pupil dilation based on its ERP and ERD data. We obtained findings qualitatively similar to the here reported between-subject analyses, see: https://doi.org/10.17605/OSF.IO/G9FQJ.

### Multimodal assessment of noradrenergic responsiveness is linked to selective attention in younger and older adults

We integrated over (pupil-associated) EEG and pupil dilation markers to derive a single, latent multimodal measure reflecting LC-NE responsiveness (see Figure 2, upper part). The proposed model fit the data well (*χ*^2^ = 6.935, *df* = 16; RMSEA = 0.0; CFI = 1.205; Brown, 2006). The variances of the latent factors differed reliably from zero in each age group (all Δ*χ*^2^ ≥ 23.845, Δ*df* = 1, all *p* < 0.001), indicating interindividual differences in NE responsiveness. Older age was associated with lower NE responsiveness scores (*rho* = –0.301; *p* = 0.006; see Figure 11 a).

**Figure 11.**
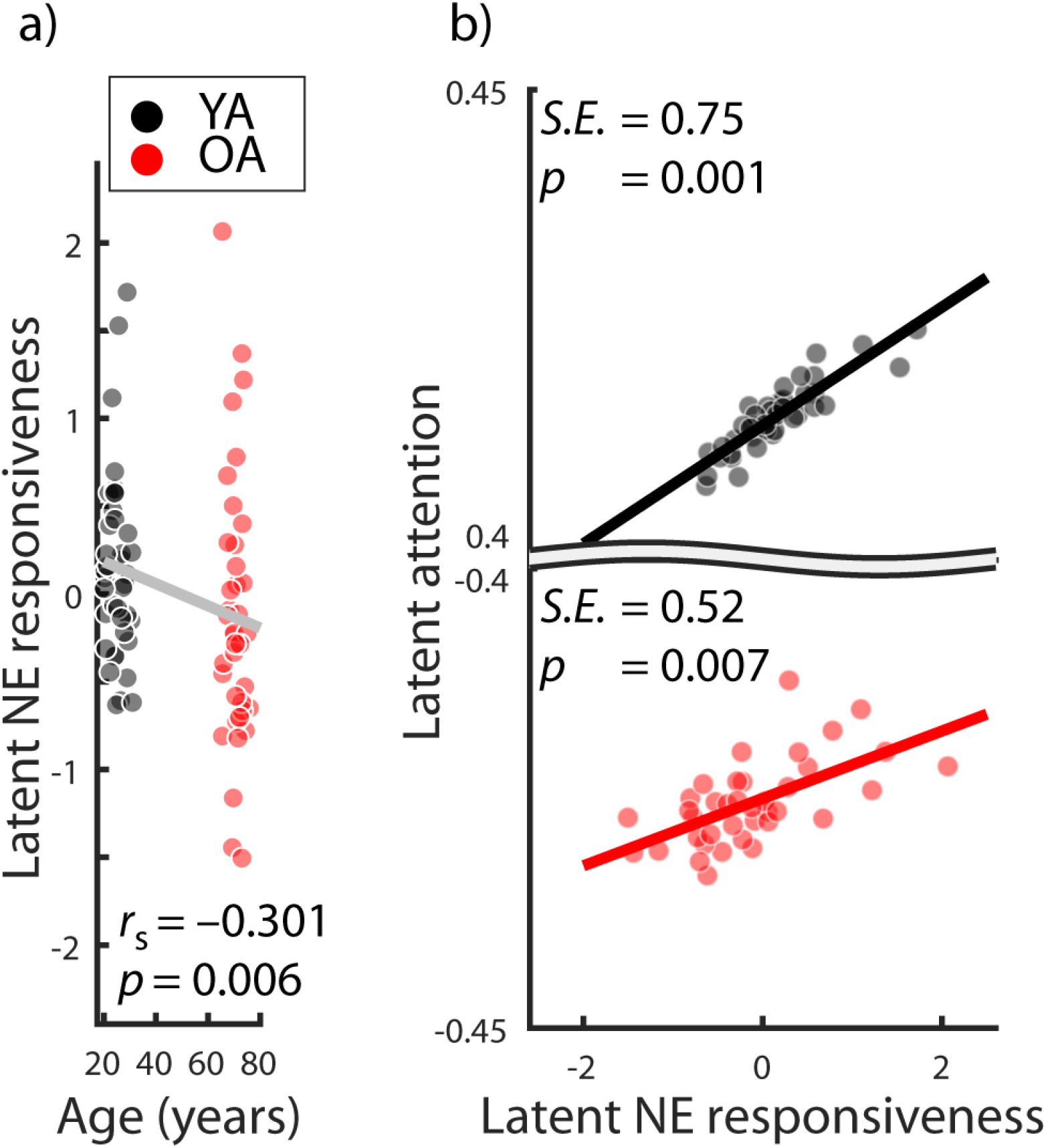
Associations between (a) age and latent norepinephrine (NE) responsiveness, evaluated using Spearman’s correlation, as well as (b) latent NE responsiveness and latent attention in younger adults (YA; black) and older adults (OA; red). Note the broken y-axis in panel (b). S.E. = standardized estimate.

To evaluate the behavioral relevance of interindividual differences in NE responsiveness in younger and older adults, we merged our general attention (see above) and NE responsiveness SEM in a unified neurocognitive model that demonstrated good fit to the data (*χ*^2^ = 45.624, *df* = 85; RMSEA = 0.0; CFI = 1.625; Brown, 2006).

Importantly, general attention was positively associated with latent NE responsiveness scores in both younger and older adults (YA: Δ*χ*^2^ = 10.323, Δ*df* = 1, *p* **=** 0.001, standardized estimate = 0.75; OA: Δ*χ*^2^ = 7.262, Δ*df* = 1, *p* **=** 0.007, standardized estimate = 0.52; see Figure 6b). The strength of the NE–attention association did not differ reliably between age groups (Δ*χ*^2^ = 0.003, Δ*df* = 1, *p* **=** 0.954). This indicates that, in face of declining selective attention in aging, a responsive NE system was linked to preserved cognitive abilities (Nyberg et al., 2012). Notably, qualitatively similar results were obtained when we analyzing composite scores of noradrenergic responsiveness and selective attention (i.e., without relying on a structural equation model; see: https://doi.org/10.17605/OSF.IO/G9FQJ).

## Discussion

Animal studies suggest that attention deficits in aging are linked to altered central noradrenergic activity (Arnsten and Goldman-Rakic, 1985; Ramos et al., 2006). In vivo research in aging humans, however, was long hampered by methodological challenges in the reliable assessment of LC-NE activity (Astafiev et al., 2010). Here we build on recent reports that pupil dilation (Joshi et al., 2016; Reimer et al., 2016; Breton-Provencher and Sur, 2019; Deitcher et al., 2019; Zerbi et al., 2019) and certain event-related EEG components (Harris and Thiele, 2011; Marzo et al., 2014; Neves et al., 2018; Vazey et al., 2018) are valid, non-invasive proxies for noradrenergic activity. In particular, we made use of LC-NE’s well-established role in fear processing (Rasmussen and Jacobs, 1986; Szabadi, 2012; Uematsu et al., 2017) to experimentally test the responsiveness of the central noradrenergic system while recording pupil dilation and the EEG. In addition, we applied a multimodal assessment to probe general attention performance in samples of healthy younger and older adults. Our findings demonstrate impaired attention in aging across multiple tasks. Moreover, older age was associated with a reduced NE responsiveness as indexed by pupil dilation and EEG. Crucially, within both younger and older adults individual differences in attention were positively related to the responsiveness of the noradrenergic system.

On the behavioral level, both younger and older adults demonstrated successful auditory selective attention in a dichotic listening task (Hugdahl et al., 2009). That is, participants in both age groups were able to adapt their attentional focus according to changing demands. However, in line with earlier reports (Passow et al., 2012, 2014; Dahl et al., 2019a), older adults showed impaired attention performance in the dichotic listening task and beyond that, across a variety of alternative attention tasks (Kennedy & Mather, 2019).

On the physiological level, we observed a multimodal response to the arousal manipulation during fear (re)conditioning in younger and older adults. In particular, compared to perceptually matched control stimuli (CS–), conditioned stimuli (CS+) elicited a sustained dilation of the pupil, as previously reported (e.g., Lee et al., 2018). In line with recent animal work linking LC activity to pupil dilation, we thus conclude that our manipulation successfully activated the LC-NE system in younger and older adults (Rasmussen and Jacobs, 1986; Szabadi, 2012; Uematsu et al., 2017; Deitcher et al., 2019).

However, using non-invasive measures we cannot rule out that other arousal-related neuromodulatory systems also influenced pupil diameter ((Reimer et al., 2016).

Conditioned stimuli (CS+) further gave rise to two sustained centro-parietal event-related EEG components: First, an anticipatory slow wave and second, a late parietal potential that occurred before and after onset of the reinforcement (US), respectively. A comparable slow wave (Stimulus-Preceding Negativity; SPN), has been observed in response to cues (S1) that prepared participants for the occurrence of following arousing or behaviorally relevant stimuli (S2; cf. Breska and Deouell, 2017; for a review see van Boxtel and Böcker, 2004). Concerning its functional relevance, the SPN has been suggested as marker of anticipatory processes that adjust the excitability of cortical networks to facilitate subsequent processing (of S2; Birbaumer, Elbert, Canavan, & Rockstroh, 1990). Similarly, Brunia (Brunia, 1993) proposed the SPN as index of regionally-targeted changes in cortical excitability that are produced via cortico-thalamic interactions. Interestingly, various peripheral correlates of noradrenergic activation (e.g., skin conductance, heart rate, cf. Szabadi, 2013) have been observed concomitant with the SPN (van Boxtel and Böcker, 2004; Poli et al., 2007). Taken together, larger SPN following fear conditioned stimuli (CS+ vs. CS–) point to a heightened anticipatory attention deployment in arousing situations in younger and older adults. After presentation of the reinforcement (US), conditioned stimuli (CS+) were associated with a second sustained parietal event-related component (Late Parietal Potential; LPP). Previous studies have observed the LPP during fear conditioning (Bacigalupo and Luck, 2018) and suggested it as index of facilitated attention allocation to arousing stimuli (for review, see Schupp et al., 2006). Furthermore, the LPP has been linked with peripheral markers of noradrenergic activity (e.g., skin conductance response; cf. Szabadi, 2013) and subjective arousal ratings. In line with previous work, we interpret the LPP as reflecting elevated selective attention during arousing conditions in younger and older adults.

In addition to ERP, fear conditioned stimuli (CS+) produced pronounced changes in rhythmic neural activity within younger and older adults. We observed an anticipatory, long-lasting desynchronization in low EEG frequencies (ERD) with strongest magnitude at parieto-occipital electrodes (van Boxtel & Böcker, 2004). Increased activity in neuromodulatory nuclei like the LC causes global cortical desynchronization (i.e., cortical state changes; McCormick et al., 1991; Marzo et al., 2014; Neves et al., 2018). Of note, the neural patterns associated with cortical state changes and selective attention are highly similar (Harris and Thiele, 2011; Thiele and Bellgrove, 2018). In particular, the global, LC-NE mediated cortical desynchronization may achieve the spatial precision necessary to selectively process attended stimuli in interaction with glutamate (Harris and Thiele, 2011; Mather et al., 2016). A wide range of human EEG studies established desynchronization in the alpha–beta range as marker of decreased cortical inhibition that allows for facilitated information processing (Hanslmayr, Staudigl, & Fellner, 2012; Jensen & Mazaheri, 2010; Klimesch, Sauseng, & Hanslmayr, 2007). Employing a similar fear conditioning procedure in rats, Headley and Weinberger (2013) demonstrated that the CS+ induced decrease in low frequencies is accompanied by a strong increase in high frequency multiunit activity (gamma; 40–120 Hz), indicating facilitated feed-forward processing (Fries, 2005, 2015). In line with earlier work, the pronounced low frequency desynchronization in response to conditioned stimuli (and US) suggests a transition towards a more activated cortical state, including increased cortical excitability and attention deployment.

Most of the arousal responses observed during (re)conditioning (i.e., ERPR, ERP; ERD) persisted in both age groups and were reinstated during the dichotic listening task in the absence of reinforcements (US). As a notable exception, in older adults, pupil reinstatement did not reach significance on a group level, potentially reflecting age-related difficulties in triggering and maintaining self-initiated processing (Lindenberger and Mayr, 2014). That is, during re/conditioning repeated external reminders (i.e., US) may have supported older adults and thus obscured age differences in pupil responses. In contrast, the lack of this external support during the dichotic listening task may have specifically affected older adults and revealed underlying age-related differences in fear conditionability (LaBar et al., 2004) and the central noradrenergic system (Betts et al., 2017; Dahl et al., 2019b; Liu et al., 2019). In line with this notion, older adults demonstrated reliable modulation of pupil dilation during phases of high external support (i.e., encoding of series of visually presented digits) but no significant modulation during phases requiring more self-initiated processing (i.e., cued recall; see Figure 1 and 3 in Van Gerven et al., 2004).

Crucially, however, in both age groups, individual differences in pupil reinstatement were linked to EEG correlates of the arousal response (i.e. SPN-ERP, ERD), suggesting a common underlying factor. The association between pupil dilation, i.e. our index of LC activity, and EEG responses is in line with optogenetic and pharmacological animal studies (Berridge and Waterhouse, 2003; Vazey et al., 2018). In particular, Vazey and colleagues (2018) demonstrated that LC photoactivation produced a positive cortical ERP ∼140–400 ms after LC stimulation in the absence of sensory input (cf. Nieuwenhuis et al., 2005). Both the parietal topography as well the time course of the observed pupil-associated ERP cluster overlap with such a LC-induced parietal positivity (i.e., 108–214 ms after t = 2; [i.e., the onset of the reinforcement during (re)conditioning]). Further, pharmacological animal studies causally implicate LC activity in the modulation of cortical and behavioral states (for a review see Berridge and Waterhouse, 2003). This effect is presumably mediated via NE’s action in the thalamus and an activation of the basal forebrain (Buzsáki et al., 1988, 1991; McCormick, 1989; McCormick et al., 1991). Behaviorally significant environmental stimuli elicit a reflexive (re)orienting of attention (orienting response) that is tightly linked to LC activity (Bouret and Sara, 2005; Sara and Bouret, 2012). Remarkably, the orienting response is always accompanied by EEG desynchronization and pupil dilation (Sara and Bouret, 2012), supporting a common dependency on LC activity.

We thus integrated over (pupil-associated) EEG and pupil dilation markers to derive a single, multimodal measure reflecting LC-NE responsiveness. We observed a lower NE responsiveness with older age which complements previous reports of structural age differences in the LC(see: https://doi.org/10.17605/OSF.IO/G9FQJ and: Betts et al., 2017; Dahl et al., 2019b; Liu et al., 2019) and age differences in LC functional connectivity (Lee et al., 2018). Crucially, within both younger and older adults, a higher noradrenergic responsiveness was associated with better selective attention performance (cf. Arnsten & Goldman-Rakic, 1985). That is, in the face of declining selective attention in aging, a responsive NE system was linked to preserved cognitive abilities (Nyberg et al., 2012).

Notably, however, as our study reports correlational data, we cannot exclude that better attentional abilities may have facilitated a preferential processing of conditioned stimuli which in turn may have led to increased noradrenergic drive. In addition, while previous research demonstrated a link between LC activity and pupil dilation (LC → pupil), increases in dilation do not necessarily imply that (only) changes in LC-NE activity have occurred. In this study we applied an experimental manipulation, fear conditioning, that reliably drives LC-NE activity, as indicated by markers of neuronal activity in animals (e.g., LC spiking activity: Rasmussen and Jacobs, 1986, Ca2+ responses in LC axons: Deitcher et al., 2019, and c-Fos: Uematsu et al., 2017; cf. Szabadi, 2012). Combining a manipulation that elicits LC activation (fear conditioning) with a non-invasive marker sensitive to LC activation (pupil dilation), we believe that our findings are at least partly attributable to the effects of NE. This argument is supported by our finding that pupil dilation was associated with electrophysiological indices that have also been linked to NE activity in invasive animal studies (i.e., the P300 ERP: Nieuwenhuis et al., 2005; Vazey and Aston-Jones, 2014; and low frequency desynchronization: McCormick et al., 1991; Marzo et al., 2014; Neves et al., 2018). Targeting the noradrenergic system from multiple angles we hope to narrow down our conclusions. Finally, using LC-MRI recordings (see https://doi.org/10.17605/OSF.IO/G9FQJ), we observed a reliable association between our NE responsiveness factor and peak LC-MRI contrast.

To conclude, we used non-invasive in-vivo markers of noradrenergic activity (Joshi et al., 2016; Reimer et al., 2016; Neves et al., 2018; Vazey et al., 2018) to uncover age differences in NE responsiveness. Importantly, structural equation modeling revealed reliable positive associations between NE responsiveness and attention in both younger and older adults. Our findings link animal and human studies on the neural underpinning of selective attention in aging (Arnsten & Li, 2005) and underscore the importance of the LC-NE system in late life cognition (Wilson et al., 2013; Mather and Harley, 2016).

## Acknowledgments

MW-B received support from the German Research Foundation (DFG, WE 4269/5-1) and the Jacobs Foundation (Early Career Research Fellowship 2017–2019). MCS was supported by the MINERVA program of the Max Planck Society and the German Research Foundation (DFG, BR 4918/2-1). MJD is a fellow of the International Max Planck Research School on the Life Course (LIFE; http://www.imprs-life.mpg.de/en). MJD is recipient of a stipend from the Sonnenfeld-Foundation (http://www.sonnenfeld-stiftung.de/en/). MM’s work was supported by an Alexander von Humboldt fellowship, a Max Planck Sabbatical Award, and by National Institutes of Health grant R01AG025340. We thank all participants and student assistants involved in the study and in particular I. Boux, Y-J Yi, M. Lawes, and N. Doehring for valuable help during data collection and preprocessing.

